# Autophagy functions in lung macrophages and dendritic cells to regulate allergen-dependent inflammatory responses

**DOI:** 10.1101/2023.03.16.533006

**Authors:** Neha Dubey, Reilly Woodson, Skyler V. Hendrix, Anne L. Rosen, Rachel L. Kinsella, Samuel R. McKee, Marick Starick, Nicole Rivera-Espinal, Sumanta K. Naik, Asya Smirnov, Darren Kreamalmeyer, Andrew L. Kau, Christina L. Stallings

## Abstract

Asthma affects 260 million people worldwide, with severe asthma cases that are associated with T_H_17/T_H_1 responses and neutrophil dominated inflammation being the most difficult to treat due to corticosteroid insensitivity. Single nucleotide polymorphisms in the *ATG5* gene, which encodes for a protein required for the cellular recycling process of autophagy, are associated with higher risk for developing severe asthma. However, the role for ATG5 during allergic inflammation remains mostly unknown. We have identified an autophagy-dependent role for ATG5 in lung macrophages and dendritic cells (DCs) for suppressing T_H_17 responses and neutrophil accumulation in house dust mite (HDM)-challenged mice, a T_H_17/T_H_1 dominated model for allergic airway inflammation due to contamination of the HDM with lipopolysaccharide. In contrast, autophagy was required to promote eosinophil accumulation in the T_H_2-dominated ovalbumin model of allergic airway inflammation, supporting a model where autophagy functions in lung macrophages and DCs to suppress T_H_17 responses and promote T_H_2 responses in an allergen-dependent manner. In addition, we discover that autophagy is also required in macrophages exposed to HDM to suppress the secretion of cytokines and chemokines that would otherwise recruit neutrophils to the lungs, independent of T cell responses. Together, our data identify multiple roles for autophagy in suppressing the neutrophil accumulation in lungs that is associated with severe asthma.

## INTRODUCTION

Asthma affects nearly 260 million people worldwide[1] and involves a deleterious inflammatory response that results in immune cell infiltration in the lungs, bronchoconstriction, airway remodeling, and airway hyperresponsiveness (AHR)[2]. Asthma and allergic airway inflammation occurs in response to a wide variety of substances called allergens that induce varying levels of immune hypersensitivity[3]. The most prevalent form of asthma is associated with T_H_2 responses signaled through airway epithelial cell-derived alarmins IL-33 and IL-25, which stimulate the production of IL-4, IL-5, IL-13, and TGF-β cytokines by airway cells and an influx of eosinophils and mast cells, along with other inflammatory cells, into the lungs[4–6]. Although this form of the disease is usually less severe and responsive to conventional corticosteroid therapy[4], asthma can also present with other distinct immunological and pathological features[7–9]. In particular, 5-15% of the total asthma cases involve a T_H_17-dominated response, leading to neutrophil recruitment and accumulation in the lungs[10,11]. Increased accumulation of neutrophils is associated with severe asthma[10,11], which is clinically defined by resistance to corticosteroid therapy[8,12–14]. In addition, T_H_1 responses mediated by interferon-γ (IFN-γ) production by CD4^+^ T cells are also associated with severe asthma cases[15,16]. The association of heightened T_H_17 responses, T_H_1 responses, and neutrophil accumulation with AHR and corticosteroid resistance has been reported during severe asthma in both mice and humans[14–17].

A better understanding of the immunological processes that contribute to severe asthma is required to develop alternative therapies for when corticosteroids fail. Several studies attempting to map genetic associations with asthma have identified a link between the *ATG5* gene and susceptibility to severe asthma. A single nucleotide polymorphism (SNP) in the third intron of the *ATG5* gene (rs12212740, alleles G/A, minor variant: G) was associated with lower pre-bronchodilator forced expiratory volume (FEV1) and a risk of moderate to severe asthma in non-Hispanic Caucasian children[18]. In a separate genome wide association study, a SNP in the 5’ untranslated region (UTR)/annotated promoter region of the *ATG5* gene (rs510432, alleles G/A, minor variant: G) was associated with increased risk of asthma in children[19]. The G variant allele of SNP rs510432 results in increased promoter activity and higher expression of *ATG5* mRNA in nasal mucosal cells isolated from the children with acute asthma compared to non-acute and healthy controls. Another study reported the association of the G variant allele of SNP rs510432 with increased sputum neutrophil counts in adult asthma patients, establishing a correlation between dysregulated *ATG5* expression and neutrophil recruitment during asthma[20].

ATG5 is essential for the process of autophagy, which involves multiple protein complexes that target cytosolic components and organelles to the lysosome for degradation[21–23]. The ULK1 complex (ULK1/ULK2, ATG13, FIP200, ATG101) and PI3 kinase complex (ATG14, BECLIN1, VPS15, and VPS34) initiate autophagy by generating a phagophore from the membranes of the endoplasmic reticulum[24,25]. Elongation of the autophagosomal double membrane depends on two ubiquitin-like conjugation systems. In the first system, ATG12 is activated by ATG7, transferred to ATG10, and covalently attached to ATG5. In the second system, LC3 (microtubule-associated protein 1 light chain 3) is conjugated to phosphatidylethanolamine to generate the membrane bound form LC3-II through the actions of ATG7, ATG3, and the ATG5-ATG12/ATG16L complex[25,26]. The autophagosome membrane is then completed and targeted for fusion with the lysosome, where the autophagosome cargo are degraded. Several studies have reported that the expression of genes required for autophagy, including *ATG5*, are elevated in the lungs of asthma patients compared to non-asthmatic individuals[27–29]. Furthermore, LC3-II expression is increased in the sputum granulocytes from severe asthma patients compared to non-severe asthma patients[20,30]. Despite the genetic associations of ATG5 and autophagy with severe asthma, the precise roles for autophagy during asthma have remained elusive. Several experimental studies have implicated both deleterious and protective functions of autophagy during asthma[31], creating uncertainty in the field. There are likely multiple contributors to the discrepancies. For example, in most of the experimental studies so far, the interpretations were largely made based on pleiotropic chemical treatments that impact autophagy, among other pathways[32–38]. In addition, autophagy likely plays cell-type specific roles during exposure to allergens, which may complicate studies that use chemical treatments or globally knockdown autophagy gene expression[14].

Innate immune cells are a primary contributor to lung inflammation during asthma and ATG5 has been shown to function both dependent and independent of autophagy to regulate inflammatory responses in innate immune cells during infection[21,22,39–42]. However, studies into the roles for autophagy specifically in innate immune cells during asthma and allergic airway inflammation have been limited. In the one study that focused on ATG5 in innate immune cells, loss of *Atg5* in hematopoietic cells was sufficient to cause increased neutrophils and macrophages in bronchoalveolar lavages (BAL), increased AHR, and higher levels of IL-17 and IL-1β in response to house dust mites (HDM) challenge in mice[14]. When *Atg5*-deficient bone marrow cells were cultured in the presence of GM-CSF to generate a mixed culture of dendritic cells (DCs), macrophages, and granulocytes, and treated with HDM, the cells produced higher levels of IL-1α, IL-1β, IL-23, and caspase-1 as compared to wild type (WT) GM-CSF cultures. *Atg5*^-/-^ GM-CSF cultures also caused a higher level of IL-17 production in a co-culture with CD4^+^ T cells as compared to WT GM-CSF cultures. Transfer of *Atg5*^-/-^ GM-CSF cultured cells loaded with HDM into WT mice intratracheally resulted in increased neutrophilic inflammation and IL-17 production as compared to transfer of WT GM-CSF cultured cells, indicating that ATG5 is important to control neutrophil recruitment in response to HDM exposure. These studies support a role for ATG5 in innate immune cells to control neutrophilic inflammation during allergic challenge. However, it is still unknown which innate immune cell types specifically require ATG5 and whether its role in controlling neutrophilic inflammation during allergic challenge is dependent on autophagy, since ATG5 and other autophagy proteins have been shown to function outside of autophagy to regulate inflammation[21,23,41,42]. To address these gaps in understanding, we used genetic tools to dissect the autophagy-dependent and independent roles for ATG5 in specific innate immune cells in two allergic airway inflammation mice models. We find that autophagy is specifically required in CD11c^+^ lung macrophages and DCs to shift the balance away from T_H_17 responses and towards T_H_2 responses in an allergen-dependent manner. In addition, autophagy functions in macrophages in a T cell-independent manner to suppress pathogen associated molecular pattern (PAMP)-triggered cytokine and chemokine production that would otherwise recruit neutrophils to the lung. Thus, autophagy impacts multiple different processes within innate immune cells that are important for regulating inflammation in response to exposure to allergens.

## RESULTS

### ATG5 is required in LysM^+^ cells to promote eosinophilia and to suppress neutrophilia in airways following allergen exposure

The role of ATG5 and autophagy in specific innate immune cell types during asthma has yet to be tested since all prior work was done with general administration of pleiotropic and non-cell type specific treatments[28,32,36,37] or using cell cultures containing a mixed population of cell types[14]. To determine if ATG5 functions in innate immune cells during allergic airway inflammation, we used *Atg5^fl/fl^-LysM-Cre* mice[21,39,43,44], which specifically delete *Atg5* in monocytes, macrophages, neutrophils, and some DCs. Flow cytometric analysis of the immune cell populations in the naïve mouse lungs showed that the only cell type that was significantly different in its abundance in the *Atg5^fl/fl^-LysM-Cre* and *Atg5^fl/fl^* mice was a higher number of non-alveolar macrophages in the lungs *Atg5^fl/fl^-LysM-Cre* mice compared to the *Atg5^fl/fl^* mice (**Figure S1A**). Whereas, the numbers of neutrophils, eosinophils, alveolar macrophages, DCs, inflammatory monocytes, naïve CD4^+^ T cells (CD44^-^CD62L^+^), activated CD4^+^ T cells (CD44^+^CD62L^-^), regulatory T cells (T_regs_) (CD4^+^CD25^+^), and CD8^+^ T cells were similar between the between *Atg5^fl/fl^-LysM-Cre* and *Atg5^fl/fl^* mice (**Figure S1A**). To determine the role of ATG5 during allergic airway inflammation, we performed an ovalbumin (OVA) induced allergic airway inflammation model[45] (**Figure 1A**), which induces a T_H_2-dominanted allergic inflammatory response in lungs. We sensitized the *Atg5^fl/fl^-LysM-Cre* mice and *Atg5^fl/fl^* control mice by intraperitoneal (i.p.) administration of 50 μg purified OVA coupled with an aluminum potassium sulfate adjuvant (alum) on days 0, 7, and 14, before challenging the mice intranasally (i.n.) with 1 mg OVA on days 20, 21, and 22. Analysis of the immune cell populations in the lungs on day 23 compared to naïve mice confirmed that both eosinophils and neutrophils were recruited to the lungs in response to the OVA challenge, but the inflammatory response to OVA challenge was dominated by eosinophils in the *Atg5^fl/fl^* mice (**Figures 1B-C, S1A**), as would be expected in a T_H_2 dominant model. There was no difference in the number of neutrophils in the lungs of OVA-challenged *Atg5^fl/fl^-LysM-Cre* and *Atg5^fl/fl^* mice (**Figure 1C**). However, OVA-challenged *Atg5^fl/fl^-LysM-Cre* mice accumulated significantly lower numbers of eosinophils in the lungs compared to *Atg5^fl/fl^* mice (**Figures 1B-C**), suggesting a shift away from T_H_2 responses. OVA-challenged *Atg5^fl/fl^-LysM-Cre* mice also accumulated significantly lower numbers of non-alveolar macrophages, DCs, and CD4^+^ T cells in the lungs compared to OVA-challenged *Atg5^fl/fl^* mice (**Figure 1C**). There were no differences in the number of alveolar macrophages, inflammatory monocytes, B cells, CD8^+^ T cells, and T_regs_ in the lungs of OVA-challenged *Atg5^fl/fl^-LysM-Cre* and *Atg5^fl/fl^* mice (**Figure S1B)**. We also analyzed immune cell accumulation specifically in the airways by instilling a fluorescently labeled αCD45.2 antibody into the airways before analysis by flow cytometry. The airways of OVA-challenged *Atg5^fl/fl^-LysM-Cre* mice contained significantly fewer eosinophils and a higher number of neutrophils and B cells compared to OVA-challenged *Atg5^fl/fl^* mice (**Figures 1D, S1C)**, whereas the number of the other immune cell types analyzed were similar between OVA-challenged *Atg5^fl/fl^-LysM-Cre* and *Atg5^fl/fl^* mice (**Figure S1C**).

**Figure 1.**
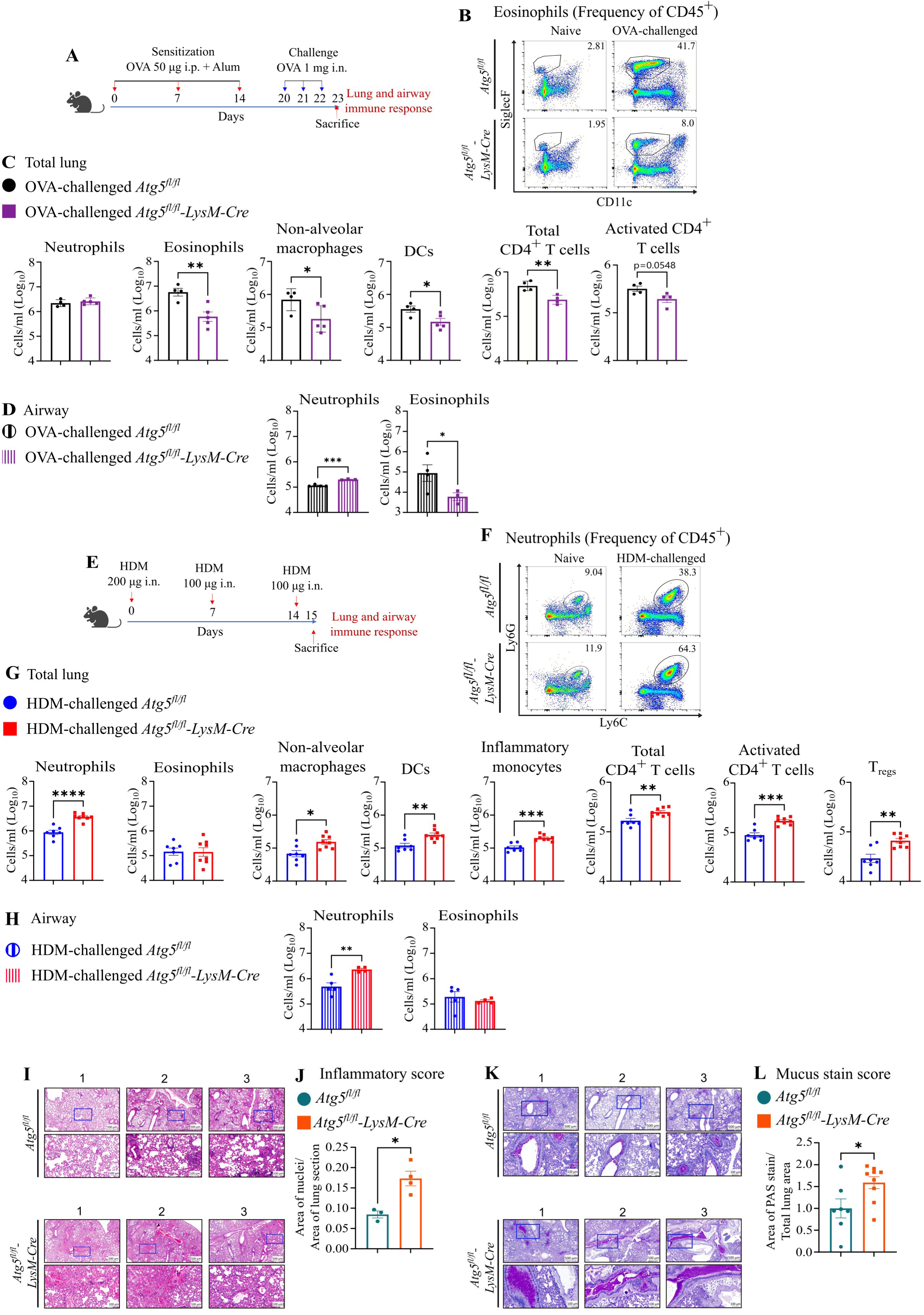
ATG5 is required in LysM^+^ cells to promote eosinophilia and to suppress neutrophilia in airways following allergen exposure. **(A)** Schematic of the OVA induced allergic airway inflammation model, made in part with BioRender. (**B**) Representative flow gating of eosinophils as a frequency of CD45^+^ cells in lungs of *Atg5^fl/fl^-LysM-Cre* and *Atg5^fl/fl^*mice. **(C)** Number of neutrophils, eosinophils, non-alveolar macrophages, DCs, total CD4^+^ T cells, and activated CD4^+^ T cells in the lungs of OVA-challenged *Atg5^fl/fl^-LysM-Cre* and *Atg5^fl/fl^* mice, represented as cells per ml of lung suspension. **(D)** Number of neutrophils and eosinophils in the airways of OVA-challenged *Atg5^fl/fl^-LysM-Cre* and *Atg5^fl/fl^*mice, represented as cells per ml of lung suspension. **(E)** Schematic of the HDM-induced allergic airway inflammation model, made in part with BioRender. (**F**) Representative flow gating of neutrophils as a frequency of CD45^+^ cells in lungs of *Atg5^fl/fl^-LysM-Cre* and *Atg5^fl/fl^* mice. **(G)** Number of neutrophils, eosinophils, non-alveolar macrophages, DCs, inflammatory monocytes, total CD4^+^ T cells, activated CD4^+^ T cells, and T_regs_ in the lungs of HDM-challenged *Atg5^fl/fl^-LysM-Cre* and *Atg5^fl/fl^* mice, represented as cells per ml of the lung suspension. **(H)** Number of neutrophils and eosinophils in the airways of the HDM-challenged *Atg5^fl/fl^-LysM-Cre* and *Atg5^fl/fl^* mice, represented as cells per ml of lung suspension. **(I,K)** Lung sections from HDM-challenged *Atg5^fl/fl^-LysM-Cre* and *Atg5^fl/fl^* mice stained with **(I)** H&E or **(K)** PAS stain. The boxed area in the top panels is zoomed in on the bottom panels. **(J,L)** Quantification of area of **(J)** inflammation based on nuclei area in H&E-stained lung histology sections and **(L)** mucus stain score calculated by analyzing the total PAS-stained area normalized to total area of lung section. Each datapoint is from an individual mouse. Data is presented on a log_10_ transformed scale except for panels J and L. The bar graphs represent the mean +/-S.E.M. Unpaired Welch’s t-test was used to determine the statistical significance of differences and only the statistically significant differences are shown. * P < 0.05, ** P < 0.01, *** P < 0.001, **** P < 0.0001.

The data from OVA-challenged *Atg5^fl/fl^-LysM-Cre* and *Atg5^fl/fl^* mice indicates that ATG5 is required in LysM^+^ cells to promote eosinophilia and suppress neutrophilia in the airways during a T_H_2-dominated allergic inflammatory response. To determine how ATG5 would impact immune responses in a T_H_17/T_H_1-dominated model for allergic airway inflammation, we used an intranasal HDM challenge model where lipopolysaccharide (LPS) contamination of the HDM has been shown to drive a primarily T_H_17/T_H_1-dominated response[46–48]. In this model, which is similar to as used in the prior study that exposed tamoxifen inducible global *Atg5^-/-^* mice to HDM and reported increased neutrophilic inflammation in the airways of *Atg5^-/-^*mice[14], HDM administered intranasally is sensed directly by innate immune cells in the lung through several pattern recognition receptors[49–51]. In addition, residual LPS contamination in the HDM is sensed by TLR4 and the DerP2 antigen in HDM mimics MD2, a TLR4 co-stimulatory molecule that confers hypersensitivity to LPS, together skewing T cell responses towards a T_H_1/T_H_17 response[46,47,52]. We challenged the *Atg5^fl/fl^-LysM-Cre* and *Atg5^fl/fl^* mice with HDM intranasally on days 0, 7 and 14 before analyzing immune cell infiltrates in the lungs and airways on day 15 by flow cytometry **(Figure 1E)**. HDM challenge resulted in the accumulation of a higher number of neutrophils than eosinophils in the lungs and airways of both *Atg5^fl/fl^-LysM-Cre* and *Atg5^fl/fl^* mice **(Figures 1F-H, S1A)**, consistent with HDM eliciting a T_H_17/T_H_1-dominated allergic airway inflammatory response. Unlike in the T_H_2-dominated OVA model, there was no difference in the number of eosinophils that accumulated in the lungs or airways of HDM-challenged *Atg5^fl/fl^-LysM-Cre* and *Atg5^fl/fl^* mice **(Figure 1G-H)**. In contrast, HDM-challenged *Atg5^fl/fl^-LysM-Cre* mice accumulated significantly more neutrophils in the lungs and airways than HDM-challenged *Atg5^fl/fl^* controls **(Figures 1F-H)**. HDM challenge of *Atg5^fl/fl^-LysM-Cre* mice also resulted in the accumulation of significantly more non-alveolar macrophages, DCs, inflammatory monocytes, activated CD4^+^ T cells, and T_regs_ in the lungs as compared to *Atg5^fl/fl^* controls **(Figures 1G)**, although the numbers of non-alveolar macrophages, DCs, inflammatory monocytes, activated CD4^+^ T cells, and T_regs_ in the airways of OVA-challenged *Atg5^fl/fl^-LysM-Cre* and *Atg5^fl/fl^* mice were similar (**Figure S1D**). There were no differences in the numbers of alveolar macrophages, B cells, naïve CD4^+^ T cells, and CD8^+^ T cells in the lungs or airways of HDM-challenged *Atg5^fl/fl^-LysM-Cre* and *Atg5^fl/fl^* mice **(Figure S1D-E).** Together, these data indicate that loss of ATG5 expression in LysM^+^ cells results in an enhanced inflammatory response to HDM challenge. Indeed, when we performed histological analysis with hematoxylin and eosin (H&E) staining of lung sections from the HDM challenged *Atg5^fl/fl^-LysM-Cre* and *Atg5^fl/fl^* mice, we observed a greater area of cellular infiltrate in the lungs of HDM-challenged *Atg5^fl/fl^-LysM-Cre* mice compared to *Atg5^fl/fl^*mice (**Figures 1I-J, S1F**), supportive of an increased inflammatory response in the lungs.

The heightened neutrophil-dominated inflammatory response in the airways of HDM-challenged *Atg5^fl/fl^-LysM-Cre* mice suggests that the mice are experiencing a more severe allergic reaction to the HDM. A characteristic feature of airway allergic reactions is the accumulation of mucus in the lungs, which increases with the severity of the allergic reaction[53]. We assessed the accumulation of neutral mucins (glycoproteins in mucus) in the airways of the HDM-challenged mice by staining with the periodic acid-schiff (PAS) dye and observed that a significantly greater area of the lung was PAS positive in HDM-challenged *Atg5^fl/fl^-LysM-Cre* mice compared to *Atg5^fl/fl^* controls (**Figures 1K-L, S1G).** Therefore, the lung pathology in HDM-challenged *Atg5^fl/fl^-LysM-Cre* mice is consistent with a more severe airway allergic reaction. These data demonstrate that ATG5 expression is required in LysM^+^ cells to suppress neutrophil-dominated inflammation in the airways of mice challenged with HDM.

### ATG5 expression in innate immune cells promotes T_H_2 responses and suppresses T_H_17 responses during allergic airway inflammation

Eosinophil and neutrophil recruitment to the airways following allergen exposure is generally associated with T_H_2 and T_H_17/T_H_1 mediated immune responses, respectively[9,46,52,54]. We investigated whether the differences in eosinophil and neutrophil recruitment to the airways of allergen challenged *Atg5^fl/fl^-LysM-Cre* and *Atg5^fl/fl^* mice reflected variances in the numbers of T_H_2 versus T_H_17 and T_H_1 cells by analyzing the expression of transcription factors that are associated with specific CD4⁺ T helper cell subsets. Specifically, we identified the CD4^+^ T cell populations expressing RORγt, GATA3, or T-bet by flow cytometry to quantify T_H_17, T_H_2, and T_H_1 cells, respectively. Similar to our prior analyses **(Figure 1C)**, the lungs of OVA-challenged *Atg5^fl/fl^-LysM-Cre* mice contained significantly fewer CD4^+^ T cells compared to OVA-challenged *Atg5^fl/fl^* mice (**Figure 2A**), where specifically the numbers of GATA3^+^ and T-bet^+^, but not RORγt^+^, CD4^+^ T cells were significantly lower in the lungs of the OVA-challenged *Atg5^fl/fl^-LysM-Cre* mice compared to OVA-challenged *Atg5^fl/fl^* mice (**Figure 2B**). Therefore, the decreased number of eosinophils recruited to the airways of OVA-challenged *Atg5^fl/fl^-LysM-Cre* mice compared to OVA-challenged *Atg5^fl/fl^* mice **(Figure 1B-D)** correlates with a decreased number of T_H_2 and T_H_1 cells in the lungs of *Atg5^fl/fl^-LysM-Cre* mice.

**Figure 2.**
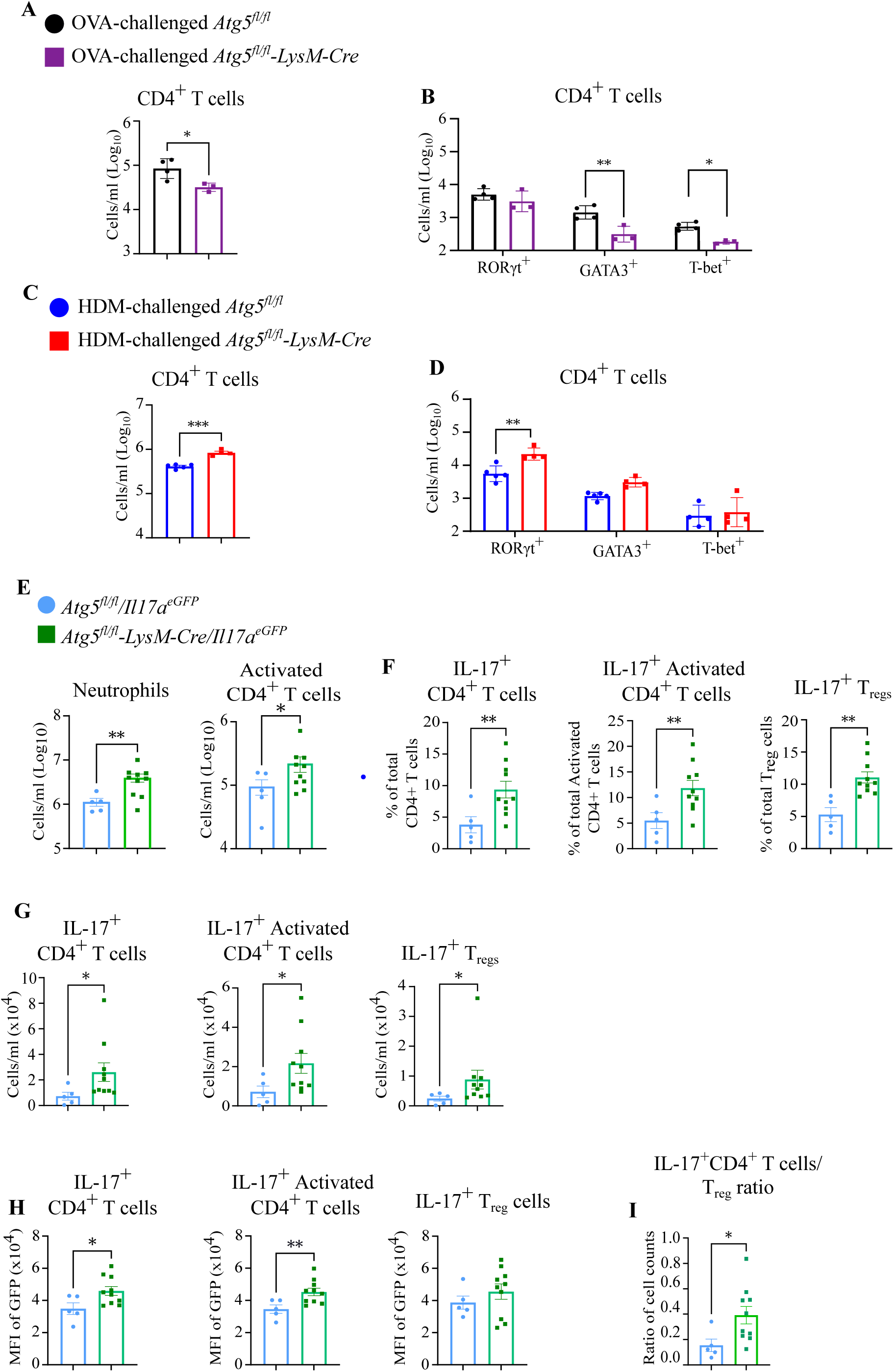
ATG5 expression in innate immune cells promotes T_H_2 responses and suppresses T_H_17 responses during allergic airway inflammation. **(A,C)** Number of total CD4^+^ T cells in the lungs of **(A)** OVA- or **(C)** HDM-challenged *Atg5^fl/fl^-LysM-Cre* and *Atg5^fl/fl^*mice, represented as cells per ml of the lung suspension. **(B,D)** Number of CD4^+^ T cells expressing RORγt, GATA3, or T-bet in the lungs of **(B)** OVA- or **(D)** HDM-challenged *Atg5^fl/fl^-LysM-Cre* and *Atg5^fl/fl^* mice, represented as cells per ml of the lung suspension. **(E)** Number of neutrophils and activated CD4^+^ T cells in the lungs of HDM challenged *Atg5^fl/fl^-LysM-Cre/Il17a*^eGFP^ and *Atg5^fl/^*^fl^/*Il17a*^eGFP^ mice, represented as cells per ml of the lung suspension. **(F)** Percentage and **(G)** number of CD4^+^ T cells, activated CD4^+^ T cells, and T_regs_ expressing IL-17A based on flow cytometry analysis of GFP expression in HDM-challenged *Atg5^fl/fl^-LysM-Cre/Il17a*^eGFP^ and *Atg5^fl/^*^fl^/*Il17a*^eGFP^ mice. **(H)** Mean fluorescence intensity (MFI) of GFP (IL-17) in CD4^+^ T cells, activated CD4^+^ T cells, and T_regs_. **(I)** Ratio of the number of IL-17^+^ CD4^+^ T cells to total T_regs_. Legend for panel **E** also applies to panels **F-I**. Each datapoint is from an individual mouse. **(A-E)** Data is presented on a log_10_ transformed scale. The bar graphs represent the mean +/-S.E.M. **(A,C, E-I)** Unpaired Welch’s t-test or **(B and D)** two-way ANOVA with Sidak’s correction was used to determine the statistical significance of differences between the groups and only the statistically significant differences are shown. * P < 0.05, ** P < 0.01.

The significantly increased numbers of CD4^+^ T cells in the lungs of HDM-challenged *Atg5^fl/fl^-LysM-Cre* mice compared to HDM-challenged *Atg5^fl/fl^* mice (**Figures 1G, 2C**) was comprised of significantly higher numbers of RORγt^+^ CD4^+^ T cells (**Figure 2D**), without differences in the numbers of GATA3^+^ or T-bet^+^ CD4^+^ T cells between the genotypes (**Figure 2D**). Thus, the increased number of neutrophils recruited to the airways of HDM-challenged *Atg5^fl/fl^-LysM-Cre* mice compared to HDM-challenged *Atg5^fl/fl^* mice **(Figure 1B-D)** correlates with an increased number of T_H_17 cells in the lungs of *Atg5^fl/fl^-LysM-Cre* mice. To specifically determine if the increased accumulation of neutrophils and RORγt^+^ CD4^+^ T cells in lungs of HDM-challenged *Atg5^fl/fl^-LysM-Cre* mice was accompanied by increased IL-17 expression from T cells, we crossed the *Atg5^fl/fl^-LysM-Cre* and *Atg5^fl/fl^* mice with an IL-17 reporter mouse[55] (*Il17a*^eGFP^, Jackson Laboratory) that expresses GFP as a proxy of IL-17, generating *Atg5^fl/fl^-LysM-Cre/Il17a*^eGFP^ and *Atg5^fl^*^/fl^/*Il17a*^eGFP^ mice. Following HDM challenge, we detected a similar increase in the abundance of neutrophils and activated CD4^+^ T cells in *Atg5^fl/fl^-LysM-Cre/Il17a*^eGFP^ mice compared to *Atg5^fl/fl^*/*Il17a*^eGFP^ controls **(Figure 2E**) as observed in the *Atg5^fl/fl^-LysM-Cre* and *Atg5^fl/^*^fl^ parent strains **(Figures 1E-F).** We monitored GFP expression from immune cells following HDM challenge of *Atg5^fl/fl^-LysM-Cre/Il17a*^eGFP^ and *Atg5^fl/fl^*/*Il17a*^eGFP^ mice using flow cytometry and found that the only cell types positive for GFP were CD4^+^ T cells. A higher percentage of total CD4^+^ T cells, activated CD4^+^ T cells, and T_regs_ expressed GFP in HDM-challenged *Atg5^fl/fl^-LysM-Cre/Il17a*^eGFP^ mice compared to *Atg5^fl/fl^*/*Il17a*^eGFP^ mice **(Figure 2F)**, which reflected a higher number of GFP^+^ activated CD4^+^ T cells and T_regs_ in the lungs of HDM-challenged *Atg5^fl/fl^-LysM-Cre/Il17a*^eGFP^ mice compared to HDM-challenged *Atg5^fl/fl^*/*Il17a*^eGFP^ controls **(Figure 2G)**. Based on mean fluorescence intensity (MFI), the activated CD4^+^ T cells in *Atg5^fl/fl^-LysM-Cre/Il17a*^eGFP^ mice also expressed more GFP and, as a proxy, IL-17 per cell than activated CD4^+^ T cells from *Atg5^fl/fl^*/*Il17a*^eGFP^ mice following HDM challenge **(Figure 2H)**. These data indicate that loss of *Atg5* in LysM^+^ cells results in increased IL-17 expression from activated CD4^+^ T cells, likely reflecting the increased number of RORγt^+^ CD4^+^ T cells. A higher T_H_17/T_reg_ ratio is also associated with increased asthma severity, increased AHR, and corticosteroid-refractory asthma[56–60]. Indeed, we observed an increased ratio of IL-17^+^ CD4^+^ T cells to T_regs_ in HDM-challenged *Atg5^fl/fl^-LysM-Cre/Il17a*^eGFP^ mice compared to HDM-challenged *Atg5^fl/fl^*/*Il17a*^eGFP^ controls **(Figure 2I)**, supporting that loss of *Atg5* in myeloid cells results in a CD4^+^ T cell imbalance that is associated with increased asthma severity[56,57,59]. Together, our data demonstrate that ATG5 is required in innate immune cells to promote T_H_2 responses in OVA-challenged mice and to suppress T_H_17 responses in HDM-challenged mice.

### Increased T_H_17 responses in HDM-challenged *Atg5^fl/fl^-LysM-Cre* mice are partially responsible for the increased neutrophil accumulation

To investigate how ATG5 regulates inflammatory responses during exposure to airway allergens, we further dissected the role for ATG5 in the HDM challenge model. Given the association of T_H_17 responses with increased neutrophil accumulation during asthma, we investigated whether the increased T_H_17 responses in the lungs of HDM-challenged *Atg5^fl/fl^-LysM-Cre* mice contributed to the accompanying increased neutrophil accumulation. To investigate the role of T cells in the neutrophil accumulation, we depleted T cells in HDM-challenged *Atg5^fl/fl^-LysM-Cre* and *Atg5^fl/fl^* mice by administering 250 µg each of αCD4 and αCD8 neutralizing antibodies intraperitoneally 2 days before (day −2) the first HDM challenge and again on days 5 and 12 post the initial HDM challenge (**Figure 3A**). Separate groups of HDM-challenged mice received 500 µg of isotype control antibodies on the same schedule to serve as controls. Flow cytometric analysis of immune cell populations in the lungs at 15 days following the first HDM dose confirmed efficient T cell depletion in the *Atg5^fl/fl^-LysM-Cre* and *Atg5^fl/fl^*mice treated with αCD4 and αCD8 antibodies (**Figure 3B**). Similar to prior experiments **(Figures 1F-G)**, HDM-challenged *Atg5^fl/fl^-LysM-Cre* mice that received the isotype control antibody accumulated higher numbers of neutrophils in the lungs compared to HDM-challenged *Atg5^fl/fl^* mice that received the isotype control (**Figure 3C)**. T cell depletion significantly decreased the number of neutrophils in the lungs of HDM-challenged *Atg5^fl/fl^-LysM-Cre* mice (**Figure 3C**), demonstrating that T cells contribute to the neutrophil accumulation in HDM-challenged *Atg5^fl/fl^-LysM-Cre* mice. However, even after T cell depletion, the neutrophil numbers were still higher in the lungs of HDM-challenged *Atg5^fl/fl^-LysM-Cre* mice than observed in the lungs of HDM-challenged *Atg5^fl/fl^* mice (**Figure 3C**). These data suggest that although T cells contribute to neutrophil accumulation in response to HDM challenge, there exists a T cell-independent process that also promotes neutrophil accumulation in the lungs of HDM-challenged *Atg5^fl/fl^-LysM-Cre* mice.

**Figure 3.**
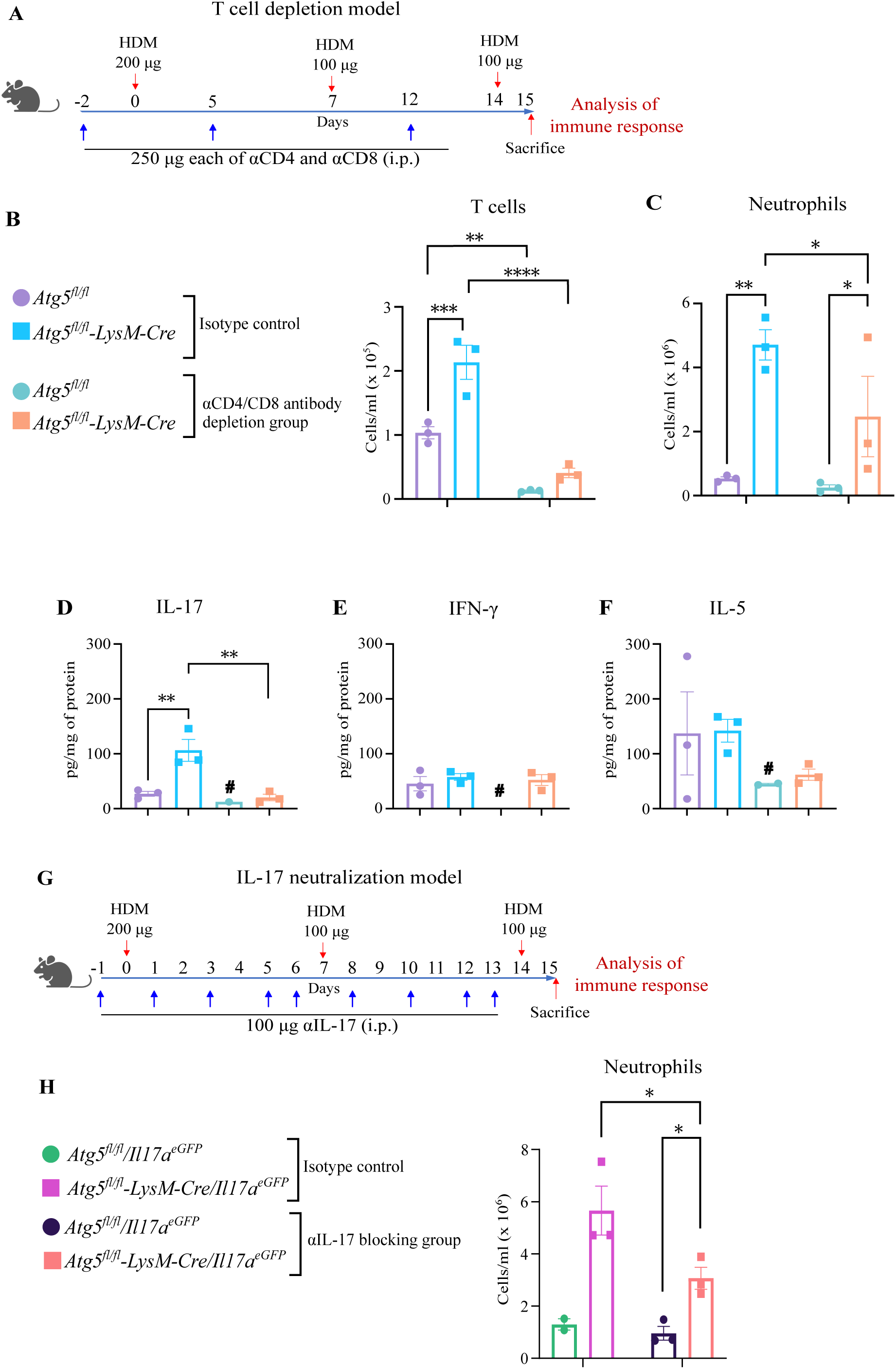
Increased T_H_17 responses in HDM-challenged *Atg5^fl/fl^-LysM-Cre* mice are partially responsible for the increased neutrophil accumulation. **(A)** Schematic of time-course of antibody mediated depletion of T cells in a murine HDM model. **(B-C)** Number of **(B)** T cells and (**C**) neutrophils in the lungs of HDM-challenged *Atg5^fl/fl^-LysM-Cre* and *Atg5^fl/fl^* mice that received either isotype control antibody or αCD4 and αCD8 antibodies, represented as cells per ml of the lung suspension. **(D-F)** Concentration of (**D**) IL-17, (**E**) IFN-γ, and (**F**) IL-5 in total lung homogenates from HDM-challenged *Atg5^fl/fl^-LysM-Cre* and *Atg5^fl/fl^* mice that received against either isotype control or αCD4 and αCD8 antibodies, based on cytokine bead array. Only the data that is above the limit of detection is plotted, where # designates conditions with samples below the limit of detection. Legend for panel B also applies to panels C-F. **(G)** Schematic of time-course of the antibody mediated blocking of IL-17 in a murine HDM model. (**H**) Number of neutrophils in the lungs of HDM-challenged *Atg5^fl/fl^-LysM-Cre* and *Atg5^fl/fl^* mice that received either isotype control antibody or αIL-17 blocking antibody. Each datapoint is from an individual mouse. The bar graphs represent the mean +/- S.E.M. Two-way ANOVA with Fisher’s LSD test was used to determine the statistical significance of differences between the groups and statistically significant differences are shown for comparisons within a genotype or within treatment groups. * P < 0.05, ** P < 0.01, *** P < 0.001, **** P < 0.0001.

To investigate the effects of T cell depletion on T_H_1, T_H_2, and T_H_17 responses, we monitored the levels of IFN-γ, IL-5, and IL-17, respectively, in the lungs of HDM-challenged *Atg5^fl/fl^-LysM-Cre* and *Atg5^fl/fl^*mice following T cell depletion versus treatment with isotype control antibodies. Within the isotype control antibody treated groups, we observed significantly higher levels of IL-17 in the lungs of HDM-challenged *Atg5^fl/fl^-LysM-Cre* mice compared to the *Atg5^fl/fl^*controls (**Figure 3D**), but no differences in the levels of IFN-γ and IL-5 (**Figure 3E, F**). This is consistent with the higher number of RORγt^+^ T_H_17 cells, but no differences in the numbers of GATA3^+^ T_H_2 or T-bet^+^ T_H_1 cells, in the lungs of HDM-challenged *Atg5^fl/fl^-LysM-Cre* mice compares to HDM-challenged *Atg5^fl/fl^* mice **(Figure 2D)**. T cell depletion significantly reduced the elevated IL-17 levels in the lungs of HDM-challenged *Atg5^fl/fl^-LysM-Cre* mice to the levels observed in the HDM-challenged *Atg5^fl/fl^* mice **(Figure 3D).** Consistent with the data from the *Atg5^fl/fl^-LysM-Cre/Il17a*^eGFP^ mice **(Figure 2),** these studies support that T cells are the primary source of IL-17 during HDM challenge and indicate that ATG5 specifically suppresses T_H_17 responses in response to HDM challenge. T cell depletion also reduced the levels of IFN-γ in the lungs of HDM-challenged *Atg5^fl/fl^* mice to below the limit of detection, but did not affect the IFN-γ levels in the HDM-challenged *Atg5^fl/fl^-LysM-Cre* mice (**Figure 3E).** These data suggest that while T cells are the major source of IFN-γ in HDM-challenged *Atg5^fl/fl^* mice, IFN-γ is produced independently of T cells in HDM-challenged *Atg5^fl/fl^-LysM-Cre*.

The T cell depletion studies indicate that enhanced T_H_17 responses contribute to the enhanced neutrophil accumulation in the lungs of HDM-challenged *Atg5^fl/fl^-LysM-Cre* mice. To directly examine the role of IL-17 signaling, we intraperitoneally administered 100 µg of an antibody that binds and blocks IL-17 1 day before the first HDM challenge (day −1) and again on days 1, 3, 5, 6, 8, 10, 12, and 13 following the initial HDM challenge (**Figure 3G).** Separate groups of HDM-challenged mice received 100 µg of isotype control antibodies on the same schedule to serve as controls. Blocking IL-17 signaling significantly reduced the number of neutrophils accumulating in the lungs of HDM-challenged *Atg5^fl/fl^-LysM-Cre* mice **(Figure 3H)**, to a similar degree as observed during T cell depletion in HDM-challenged *Atg5^fl/fl^-LysM-Cre* mice **(Figure 3C).** This supports that elevated T_H_17 responses contribute to increased neutrophil accumulation in the lungs of HDM-challenged *Atg5^fl/fl^-LysM-Cre* mice. However, similar to as observed during T cell depletion **(Figure 3C)**, the neutrophil numbers remained higher in the lungs of HDM-challenged *Atg5^fl/fl^-LysM-Cre* mice than observed in HDM-challenged *Atg5^fl/fl^* mice following αIL-17 treatment **(Figure 3H)**. Therefore, ATG5 inhibits neutrophil accumulation during HDM challenge in part through suppressing T_H_17 responses, but also through another T cell-independent mechanism.

### ATG5 is required in macrophages to control the expression of neutrophil recruitment signals during exposure to HDM

A T cell-independent inflammatory process that is negatively regulated by autophagy in innate immune cells is inflammasome activation[61–63]. Several reports have demonstrated a role for the NLRP3 (nucleotide-binding oligomerization domain-like Receptor Family Pyrin Domain Containing 3) inflammasome pathway, which catalyzes caspase-1 dependent release of pro-inflammatory cytokines IL-1β and IL-18, during severe allergic inflammation[64–67]. In addition, *Atg5*^-/-^ bone marrow cells cultured in GM-CSF produce higher levels of IL-1β than wild-type cultures following exposure to HDM *in vitro*[14]. We measured the levels of IL-1β in the lungs of HDM-challenged *Atg5^fl/fl^-LysM-Cre* and *Atg5^fl/fl^*mice and although there was a trend of higher levels of IL-1β in the HDM-challenged *Atg5^fl/fl^-LysM-Cre* mice, the IL-1β levels were not significantly different than in HDM-challenged *Atg5^fl/fl^* mice (**Figure S2A**). To directly interrogate if the increased neutrophil accumulation following HDM challenge is a consequence of inflammasome activation in the absence of autophagy in *Atg5^fl/fl^-LysM-Cre* mice, we crossed the *Atg5^fl/fl^-LysM-Cre* and *Atg5^fl/fl^* mice with *Casp1/11^-/-^* mice (Jackson Laboratory), which are deleted for the genes encoding caspase-1 and caspase-11. We challenged *Atg5^fl/fl^-LysM-Cre/Casp1/11^-/-^*and *Atg5^fl/fl^*/*Casp1/11^-/-^* mice with HDM and analyzed the lung immune cell populations. Similar to *Atg5^fl/fl^-LysM-Cre* mice, the lungs of HDM-challenged *Atg5^fl/fl^-LysM-Cre/Casp1/11^-/-^*mice accumulated more neutrophils, non-alveolar macrophages, DCs, activated CD4^+^ T cells, and T_regs_ than *Atg5^fl/fl^*/Casp1*/11^-/-^*mice, with no differences in the number of eosinophils **(Figure S2B)**. Deletion of *Casp1* and *Casp11* also did not alter the non-significant trends in IL-1β levels in the lungs of HDM-challenged *Atg5^fl/fl^-LysM-Cre* and *Atg5^fl/fl^* mice **(Figure S2C)**. These data indicate that neutrophils accumulate in the lungs of HDM-challenged *Atg5^fl/fl^-LysM-Cre* mice independent of caspase-1 and caspase-11.

To explore the T cell and inflammasome-independent mechanism by which ATG5 regulates neutrophil accumulation during HDM exposure, we examined the inflammatory signals that remained higher in the lungs of HDM-challenged *Atg5^fl/fl^-LysM-Cre* mice compared to HDM-challenged *Atg5^fl/fl^* mice during T cell depletion using a cytokine bead array **(Figures 4A, S2D)**. We found that the levels of G-CSF (granulocyte colony stimulating factor), KC (IL-8), IL-1α, IL-12p40, and MIP-1α were higher in the lungs of HDM-challenged *Atg5^fl/fl^-LysM-Cre* mice compared to HDM-challenged *Atg5^fl/fl^* mice when T cells were depleted (**Figures 4A**). G-CSF is a cytokine important for neutrophil development, KC is a neutrophil chemoattractant, and IL-1α, IL-12p40, and MIP-1α have all been associated with neutrophil recruitment during inflammation [68–71]. The higher levels of G-CSF, KC, IL-1α, IL-12p40, MIP-1α, and IFN-γ in HDM-challenged *Atg5^fl/fl^-LysM-Cre* mice depleted for T cells (**Figures 3E, 4A)** implies that the source of these inflammatory signals may be an innate immune cell.

**Figure 4.**
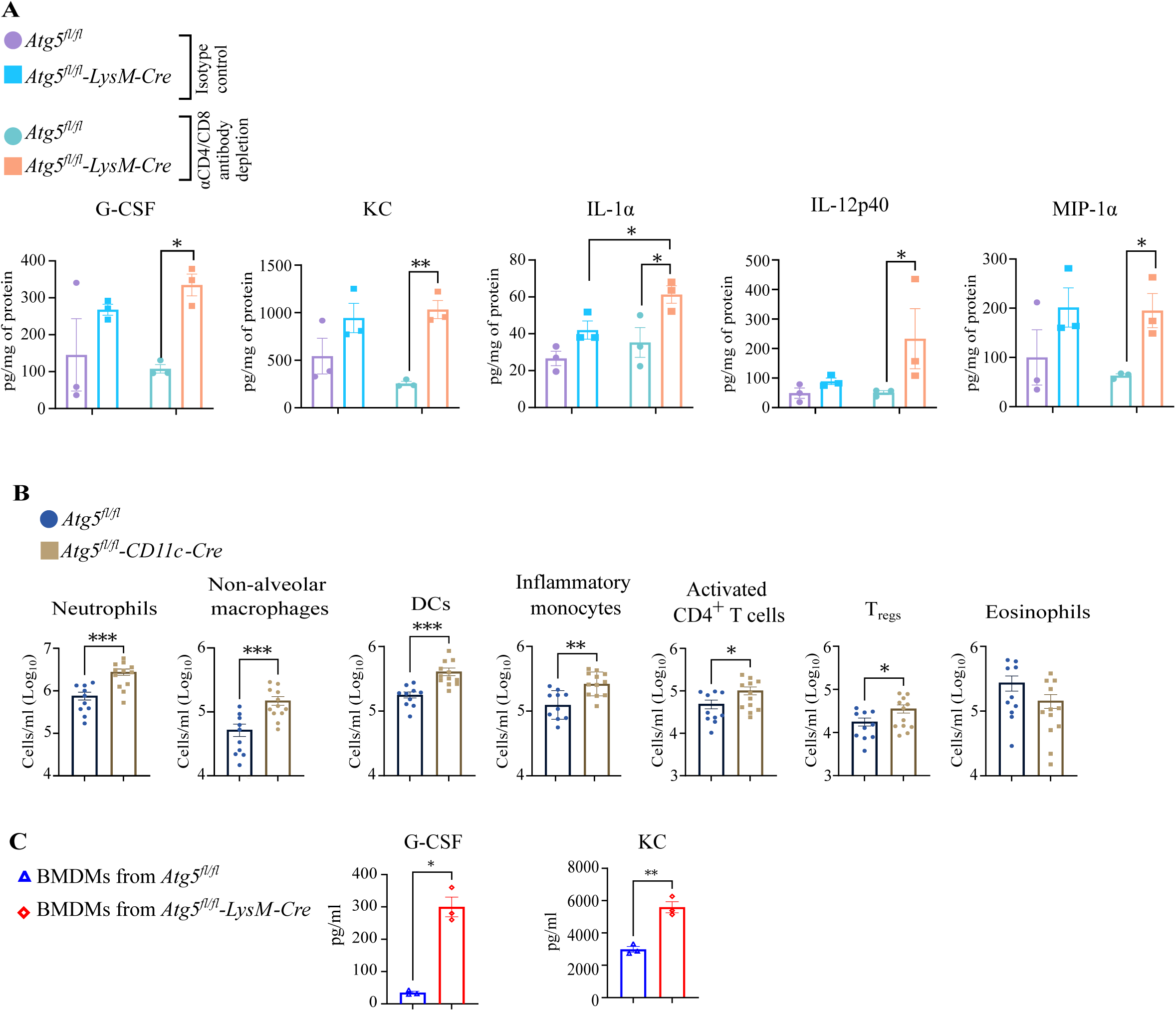
ATG5 is required in macrophages to control the expression of neutrophil recruitment signals during exposure to HDM. (**A**) Concentration of G-CSF, KC, IL-1α, IL-12p40, and MIP-1α in total lung homogenate from HDM-challenged *Atg5^fl/fl^-LysM-Cre* and *Atg5^fl/^*^fl^ mice that received against either isotype control or αCD4 and αCD8 antibodies, from cytokine bead array. Only the data that is above the limit of detection is plotted. **(B)** Number of neutrophils, non-alveolar macrophages, DCs, inflammatory monocytes, activated CD4^+^ T cells, T_regs_, and eosinophils in the lungs of HDM-challenged *Atg5^fl/fl^-CD11c-Cre* and *Atg5^fl/fl^*mice. Data is presented on a log_10_ transformed scale. **(C)** BMDMs were differentiated from bone marrow cells harvested from *Atg5^fl/fl^-LysM-Cre* and *Atg5^fl/fl^* mice and treated with or without 200 µg HDM for 24 hours. Concentration of G-CSF and KC secreted into the media by ELISA. Each datapoint is from an individual mouse. The bar graphs represent the mean +/- S.E.M. Two-way ANOVA with Fisher’s LSD test was used to determine the statistical significance of differences between the groups in panel A. Unpaired Welch’s t-test was used to determine the statistical significance of differences between the groups for panels B and C. Only the statistically significant differences are shown in the graphs. * P < 0.05 and ** P < 0.01 *** P < 0.001.

*Atg5^fl/fl^-LysM-Cre* mice delete *Atg5* in multiple different types of innate immune cells, including neutrophils, macrophages, inflammatory monocytes, and some DCs. To determine if ATG5 functioned specifically in macrophages to suppress neutrophil accumulation during HDM-challenge, we challenged *Atg5^fl/fl^-CD11c-Cre* (delete *Atg5* in lung macrophages and DCs) with HDM and monitored lung inflammatory responses. Similar to as observed in HDM-challenged *Atg5^fl/fl^-LysM-Cre* mice **(Figure 1)**, the lungs of HDM-challenged *Atg5^fl/fl^-CD11c-Cre* mice accumulated higher numbers of neutrophils, non-alveolar macrophages, DCs, inflammatory monocytes, activated CD4^+^ T cells, and T_regs_ with no difference in the number of eosinophils (**Figure 4B**) as compared to HDM-challenged *Atg5^fl/fl^* controls. In contrast, the lungs of HDM-challenged *Atg5^fl/fl^-MRP8-Cre* mice that delete *Atg5* specifically in neutrophils accumulated similar numbers of immune cells as HDM-challenged *Atg5^fl/fl^* controls (**Figures S2E**). Therefore, ATG5 functions in lung macrophages and DCs, but not neutrophils, to control neutrophil accumulation during HDM challenge.

We have previously reported that autophagy functions in macrophages independent of its role in inhibiting inflammasome activation to suppresses the production of G-CSF and KC as well as neutrophil recruitment to the lungs during *Mycobacterium tuberculosis* infection[72], however, this role for autophagy has only been studied in the context of infection. To directly test if ATG5 is required in macrophages to suppress G-CSF and KC production following the noninfectious HDM exposure, we differentiated bone marrow stem cells from *Atg5^fl/fl^-LysM-Cre* and *Atg5^fl/fl^* mice into bone marrow derived macrophages (BMDM) by culturing *ex vivo* in the presence of macrophage colony-stimulating factor (M-CSF). We treated the *Atg5^fl/fl^-LysM-Cre* and *Atg5^fl/fl^* BMDMs with 200 µg HDM for 24 hours and measured the levels of G-CSF and KC secreted into the media. The levels of G-CSF and KC produced by naïve macrophages from *Atg5^fl/fl^-LysM-Cre* and *Atg5^fl/fl^* mice was below the limit of detection. HDM treatment resulted in significantly higher levels of G-CSF and KC secreted from *Atg5^fl/fl^-LysM-Cre* macrophages compared to *Atg5^fl/fl^*macrophages **(Figures 4C)**. Therefore, ATG5 functions in macrophages to suppress G-CSF and KC production following direct sensing of PAMPs in HDM, which could otherwise promote neutrophil accumulation in lungs.

### Autophagy is required in innate immune cells to control neutrophil-dominated inflammation during HDM challenge

ATG5 has been shown to control inflammatory responses via both autophagy-dependent and autophagy-independent pathways[21,41–43,72–75]. To determine whether the regulation of neutrophil recruitment and inflammation by ATG5 in HDM-challenged mice was dependent on other autophagy proteins or represented an autophagy-independent role for ATG5, we monitored neutrophil abundance in the lungs of mice lacking expression of the essential autophagy genes *Atg16l1*, *Becn1, or Fip200* in LysM^+^ cells following HDM challenge by flow cytometry. BECLIN1 (encoded by *Becn1*) and FIP200 operate during the nucleation step of the autophagosome membrane biogenesis, whereas ATG16L1 functions with ATG5 in the phagophore elongation complex[25]. Similar to as observed in *Atg5^fl/fl^-LysM-Cre* mice, HDM-challenged *Atg16l1^fl/fl^-LysM-Cre, Becn1^fl/fl^-LysM-Cre*, and *Fip200^fl/fl^-LysM-Cre* mice accumulated more neutrophils and activated CD4^+^ T cells in their lungs compared to HDM-challenged *Atg16l1^fl/fl^*, *Becn1^fl/fl^*, and *Fip200^fl/fl^* controls, respectively **(Figures 5A-B)**. The lungs of HDM-challenged *Atg16l1^fl/fl^-LysM-Cre* and *Fip200^fl/fl^-LysM-Cre* mice also accumulated significantly higher numbers of non-alveolar macrophages, DCs, and T_regs_ in comparison to HDM-challenged *Atg16l1^fl/fl^*and *Fip200^fl/fl^* controls, respectively **(Figures 5C-E)**. The lungs of HDM-challenged *Becn1^fl/fl^-LysM-Cre* mice also accumulated a higher number of inflammatory monocytes compared to HDM-challenged *Becn1^fl/fl^* mice **(Figures 5F)**. Similar to *Atg5^fl/fl^-LysM-Cre* mice, we did not observe differences in eosinophils in the HDM-challenged *Atg16l1^fl/fl^-LysM-Cre*, *Becn1^fl/fl^-LysM-Cre*, and *Fip200^fl/fl^-LysM-Cre* mice compared to controls **(Figure 5G)**. While ATG5, ATG16L1, and BECLIN1 are essential for both canonical autophagy as well as the non-canonical autophagy processes LC3 associated phagocytosis (LAP) and LC3-dependent endocytosis (LANDO), FIP200 is dispensable for the LAP and LANDO pathways. Therefore, the requirement for ATG5, ATG16L1, BECLIN1, and FIP200 implies that canonical autophagy is necessary in LysM^+^ innate immune cells to suppress the accumulation of neutrophils in the lungs of mice following HDM challenge.

**Figure 5.**
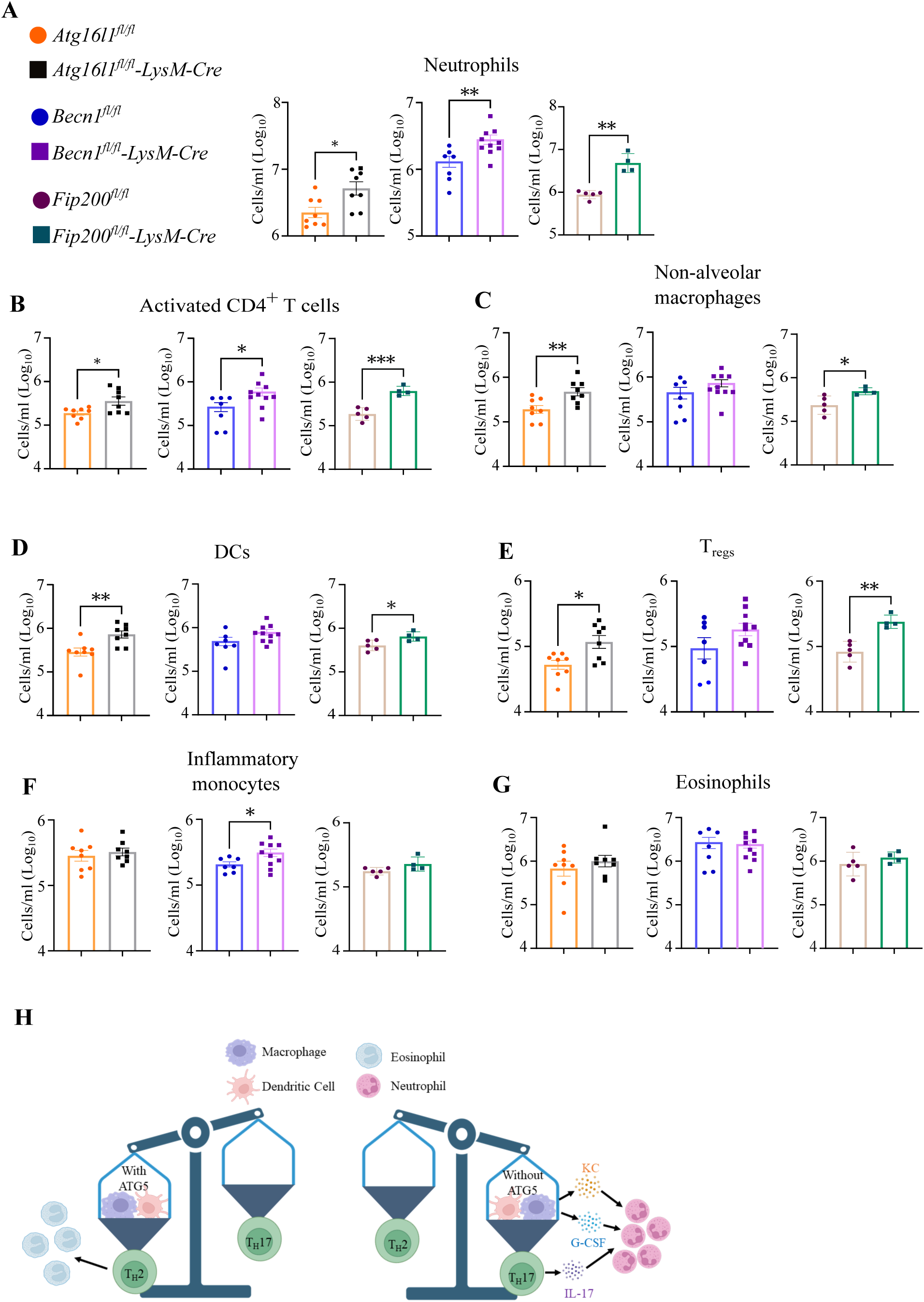
Autophagy is required in innate immune cells to control neutrophil-dominated inflammation during HDM-challenge. Number of **(A)** neutrophils (**B**) activated CD4^+^ T cells, **(C)** non-alveolar macrophages, **(D)** DCs, **(E)** T_regs_, (**F**) inflammatory monocytes, and (**G**) eosinophils in the lungs of HDM-challenged *Atg16l1^fl/fl^-LysM-Cre*, *Atg16^fl/fl^*, *Becn1^fl/fl^-LysM-Cre*, *Becn1^fl/fl^*, *Fip200^fl/fl^-LysM-Cre*, and *Fip200^fl/fl^*mice, represented as cells per ml of the lung suspension. Legends for panel A also applies to panel B-G. Each datapoint is from an individual mouse. Data is presented on a log_10_ transformed scale. The bar graphs represent the mean +/- S.E.M. Unpaired Welch’s t-test was used to determine the statistical significance of differences between the groups, and only the statistically significant differences are shown. * P < 0.05, ** P < 0.01 *** P < 0.001. (**H**) Model for the roles of ATG5 in lung macrophages and DCs during allergic airway inflammation. The image is generated in part with BioRender.

Together, our data supports a model where during airway exposure to allergens, ATG5 functions in lung macrophages and DCs to tip the balance of T cell responses to favor T_H_2 responses and suppress T_H_17 responses (**Figure 5H**). In addition, ATG5 is required in macrophages to inhibit the secretion of cytokines and chemokines that can recruit neutrophils to the lungs during allergen exposure in a T cell-independent manner. Therefore, ATG5 plays multiple roles in lung innate immune cells to limit the neutrophil recruitment and mucus accumulation that would be associated with severe asthma.

## DISCUSSION

The association of genetic polymorphisms in the *ATG5* gene with susceptibility to asthma in humans has drawn attention to the roles for autophagy in regulating inflammatory responses during allergen challenge and asthma. In the present study, we directly investigate the roles for ATG5 and autophagy in innate immune cells during asthma using a series of conditional knockout mouse lines and two murine models of allergic airway inflammation model. Our data support a model where ATG5 functions in an autophagy-dependent process in CD11c^+^ lung macrophages and/or DCs to promote T_H_2-associated eosinophil accumulation and suppress T_H_17 responses in an allergen-dependent manner. In addition to regulating T cell responses, we find that autophagy functions in macrophages exposed to HDM to suppress the secretion of cytokines and chemokines that would otherwise recruit neutrophils to the lungs, independent of T cell responses.

Our studies reveal a function for autophagy in innate immune cells that can promote T_H_2 responses while suppressing T_H_17 responses, positioning autophagy as a central regulator of T cell polarization during allergen exposure. There are multiple possible mechanisms by which autophagy could be functioning in CD11c^+^ DCs and lung macrophages to impact T cell polarization during HDM challenge. Both DCs and lung macrophages are antigen-presenting cells (APCs) that could serve to activate allergen specific CD4^+^ T cells during asthma[76–78]. Autophagy plays a central role in transferring antigen to vesicles that contain MHC class II molecules for antigen presentation to CD4^+^ T cells[79,80]. In addition to direct roles in MHC class II antigen presentation, autophagy also regulates the production of cytokines that can impact the polarization of T cells during activation. Loss of autophagy can result in increased production of IL-23 by macrophages and DCs[81], and IL-23 can induce the differentiation of T cells into T_H_17 cells[82,83]. Loss of autophagy in macrophages also promotes pro-inflammatory M1-like macrophage polarization in inflammatory conditions such as aging-associated hepatic injury and obesity[73,84], and macrophage polarization can also impact T cell responses. M1-like macrophages produce T_H_1 associated cytokines (TNF-α, IL-6, IL-1β, and IL-23) and induce a T_H_1/T_H_17 type inflammatory response, while M2-like macrophages produce type 2 cytokines (IL-5 and IL-13), activating a T_H_2 responses[85]. M1-like macrophages also express higher levels of the transcription factor IRF5, which was demonstrated to drive T_H_1/T_H_17 responses in culture[86] and deletion of *Irf5* in mice causes a reduction in IFN-γ, IL-17, and neutrophil recruitment in HDM-challenged mice[87]. Therefore, autophagy in macrophages and DCs can drive differential T cell responses via antigen presentation and the production of inflammatory signals.

Loss of autophagy in lung macrophages and DCs also resulted in the accumulation of T_regs_ in the lungs following HDM challenge. Although T_regs_ are well-known to suppress the inflammatory responses during asthma[88–91], T_reg_ accumulation is also reported to be increased during severe asthma[92,93]. Transfer of purified T_H_17 cells promoted the expansion of T_regs_ in mice[94]. Therefore, it is possible that the increased T_H_17 responses directly contribute to the increase in T_regs_ during HDM challenge of *Atg5^fl/fl^-LysM-Cre* mice, which may serve as a countermeasure to dampen the increased inflammatory response. Loss of autophagy in DCs is also associated with dysregulated T_reg_ function in mice due to an accumulation of ADAM10 metalloprotease on the surface of DCs in the absence of autophagy[95]. Accumulation of ADAM10 leads to cleavage of ICOS-L, which is required for the efficient maturation of T_regs_ in steady state. Moreover, in the absence of proper maturation, FOXP3^+^CD25^+^ T_regs_ transition to IL-17^+^ CD4^+^ T cells[95], which may explain the increased IL-17 expression in T_regs_ that we observed in HDM-challenged *Atg5^fl/fl^-LysM-Cre* mice.

In addition to its role in regulating T cell polarization during allergen exposure, we find that autophagy also functions in macrophages independent of T cells to suppress the production of IFN-γ, G-CSF, and KC in response to HDM exposure. High levels of IFN-γ, G-CSF, and IL-8 (KC) have all been associated with neutrophilic inflammation, asthma progression, and severe asthma in humans[15,96,97]. Autophagy has been reported to serve a similar role in lung macrophages during *M. tuberculosis* infection[72], however, this was assumed to be a response to the active infection. Our studies here show that autophagy more generally regulates the production of neutrophil recruitment signals from macrophages during exposure to PAMPs. This is supported by prior studies that autophagy-deficient BMDMs upregulated pro-inflammatory cytokine production in response to LPS and IFN-γ[98]. HDM is a complex mix of bioactive molecules and immune stimulatory molecules (including proteases and other allergens, β-glucans, and chitin exoskeleton) that can directly be sensed by innate immune cells through several pattern recognition receptors (TLR and CLR), NOD like receptors, and protease activated receptors[49–51]. In addition, the residual LPS present in HDM is sensed by TLR4, where increased doses of LPS shift the eosinophilic lung inflammation induced by LPS-free HDM to neutrophil-dominated inflammation[46,48]. Indeed, mice deficient in *Myd88* and *Tlr4* were found to be protected from HDM induced allergic response[99]. ATG5 has also been reported to directly interact with and regulate MyD88 in mouse embryonic fibroblast (MEF) cells by preventing the formation of MyD88 condensates that result in increased MyD88-TRAF6-NFκB signaling complex activation[100]. In addition, autophagy suppresses pro-inflammatory responses in macrophages through mitophagy[101–104], the clearance of damage-associated molecular patterns (DAMPs) and PAMPs[105–107], and ER-phagy[108,109]. These roles for autophagy could also impact neutrophil recruitment and accumulation in the lungs during HDM challenge. The increased sensitization to PAMP signaling in the absence of autophagy could also be contributing to the enhanced T_H_17 responses where autophagy promotes T_H_17 responses through increased IL-23 secretion in response to TLR agonists[81]. Given the similarities between the roles for autophagy during *M. tuberculosis* infection and HDM challenge, it is also likely that autophagy also regulates inflammation in a similar manner during other pulmonary inflammatory diseases. In addition, there is evidence that autophagy may play similar roles in other tissues where autophagy in myeloid cells alleviated intestinal inflammation and neutrophil accumulation during macrophage activation syndrome and adherent-invasive *Escherichia coli* (AIEC) infection in mice[98]. These findings highlight autophagy as a central regulator of inflammatory responses from innate and adaptive immune cells in multiple tissues and diseases.

Human asthma is a heterogeneous condition characterized by multiple endotypes with distinct cellular signatures[110]. Patients with T_H_2-high versus T_H_2-low endotypes respond differently to corticosteroid treatments, which is associated with distinct and mixed granulocytic phenotypes. In addition, there exists incredible heterogeneity in what environmental trigger causes asthma in humans, which can also include viral infections. In our mouse models, we examined one example each of T_H_2-high and T_H_2-low allergic airway responses. Using these models we have identified a protective role for autophagy in controlling neutrophil accumulation during asthma, suggesting that inducing autophagy may hold promise as a therapeutic for treating severe asthma cases. Although this provides a starting point for understanding the role of autophagy in innate immune cells during allergic airway inflammation, the application of these findings to the spectrum of diversity observed in asthma cases in humans will still require expanding the complexity of the system, a challenge for all animal models of human disease. In addition, the therapeutic potential for targeting autophagy is currently limited by the absence of autophagy-specific inducers and the multitude of diverse roles for canonical autophagy in eukaryotic cells. Therefore, the translation of these findings to treatments will benefit from future studies into the effectors that function downstream of autophagy to govern the immune responses during airway exposure to allergens.

## METHODS

### Mouse strains

All flox mice (*Atg5^fl/fl^, Becn1^fl/fl^, Atg16l1^fl/fl^*) used in this study have been described previously[39,44,111,112] and are maintained in an enhanced barrier facility. LysM-Cre (Jax #004781), CD11c-Cre (Jax #007567), and MRP8-Cre (Jax #021614) mice from the Jackson Laboratory were crossed to specific flox mice, respectively. *Atg5^fl/fl^-LysM-Cre* and *Atg5^fl/fl^* mice were crossed with *Casp1/11*^-/-^ (Jax #016621) germline deletion mice to generate *Atg5^fl/fl^-LysM-Cre/Casp1/11^-/-^* and *Atg5^fl/fl^/Casp1/11^-/-^* mice. *Atg5^fl/fl^-LysM-Cre* and *Atg5^fl/fl^* mice were crossed with *Il17a*^eGFP^ (Jax #018472) mice to generate *Atg5^fl/fl^-LysM-Cre/Il17a*^eGFP^ and *Atg5^fl/^*^fl^/*Il17a*^eGFP^ mice. Adult male and female mice (aged 8–12 weeks) were used for the experiments. All experiments were performed with respective littermate flox controls. All mice have been backcrossed to C57BL/6J background. The mice were housed and bred at Washington University in St. Louis in specific pathogen-free conditions in accordance with federal and university guidelines. All procedures involving animals were conducted following National Institute of Health guidelines and protocols were approved by the Institutional Animal Care and Use Committee of Washington University in St. Louis. Animals were euthanized by CO_2_ asphyxiation, approved by Euthanasia of the American Veterinary Association.

### Allergic airway inflammation models

Mice were used in an OVA model of airway allergic inflammation as described before, with few modifications[45]. 8-12 week old sex matched mice were sensitized with 50 µg OVA (Sigma, A5503-5G) complexed with aluminum hydroxide and magnesium hydroxide suspension (Imject™ Alum Adjuvant, Thermo scientific, 77161) on day 0, 4 and 14 intraperitoneally in 200 µl 1X PBS. Mice were challenged with 1 mg OVA in PBS in 25 µl intranasally on days 20, 21, and 22. Mice were sacrificed on day 23, and the whole lung (without the post-caval lobe) was processed for flow cytometry analysis. For the HDM model of allergic airway inflammation, 8-12 week-old sex matched mice were challenged with HDM as described previously[14]. Briefly, mice were challenged intranasally with HDM (Greer Laboratories, NC9756554) in PBS (Gibco, 10010023). The first primary dose of HDM (200 µg in 40 µl PBS) was administered on day 0 and two secondary (100 µg in 20 µl PBS) doses were administered on days 7 and 14. Mice were sacrificed on day 15 and the whole lung except the post-caval lobe was processed for the flow cytometry experiments.

### Flow cytometry

Mouse lungs were excised after PBS perfusion, placed in DMEM (Gibco, 12100046), minced finely, and digested at 37⁰C for an hour with mechanical disruption with a stir bar and enzymatic digestion with collagenase D (Roche, 11088875103) and DNase I (Sigma, D4527). Digested tissue was treated with ACK buffer (Life technologies, A1049201) to remove red blood cells and were passed through a 70 μm cell strainer to generate single cell suspension in 1 ml of semi-facs buffer (1X PBS pH 7.4 supplemented with 2% FBS and 2 mM EDTA). 100 μl of cells were stained with Zombie-NIR before staining with the antibody cocktail mix. Cells were resuspended in semi-facs buffer in the presence of Fc receptor blocking antibody (BioLegend, 101302) and stained with antibodies at a 1:200 dilution with the myeloid or lymphoid antibody cocktails. The innate immune cell staining antibody cocktail included the following antibodies: Ly6G-BV421 (Biolegend, 1A8), SiglecF-BV480 (Biolegend, E50-2440(RUO), CD45-BV570 (Biolegend, 30-F11), CD11b-BV650 (Biolegend, M1/70), CD103-BV750 (Biolegend, M290), Ly6C-BV785 (Biolegend, HK1.4), MHCII-SparkBlue550 (Biolegend, M5/114.15.2), CD64-PerCP-eFluor710 (Biolegend, X54-5/7.1), MerTK-PE-Cy7 (Biolegend, DS5MMER), CD206-eF660 (AF647) (eBiosciences, C068C2), CD11c-APC/Fire750 (Biolegend, N418). The lymphoid cell staining antibody cocktail included the following antibodies: CD8α-Pacific blue (Biolegend, 53-6.7), CD44-BV510 (Biolegend, IM7), CD45-BV570 (Biolegend, 30-F11), CD19-BV605 (Biolegend, 6D5), CD11b-BV650 (Biolegend, M1/70), NK1.1-BV785 (Biolegend, PK136), TCRβ-FITC (Biolegend, H57-597), NKp46-PerCP-eFluor710 (eBiosciences, 29A1.4), CD62L-PE/Cy5 (Biolegend, MEL-14), CD25-PE-Cy7 (Biolegend, PC61), and CD4-APC (Biolegend, 100516). For the intracellular staining, surface-stained cells were fixed with 4% PFA before proceeding for intracellular staining for the transcription factors. Cells were permeabilized with 1X FoxP3 permeabilization buffer (Biolegend, 421403) and the intracellular staining is performed for the transcription factors using fluorophore conjugated antibodies for RORγt-APC (eBiosciences, B2D), GATA3-APC (Biolegend, 16E10A23), and T-bet-APC (Biolegend, 4B10), in separate panels, as per Biolegend’s instructions. To label immune cells in the airways before performing flow cytometry analysis, mice were anesthetized with a ketamine-xylazine cocktail injected intraperitoneally. CD45.2-FITC (Biolegend, 104) was administered intratracheally in the anesthetized mice to allow for airway specific cell staining in the live mice for 30 minutes. Mice were then sacrificed, and total lungs were collected and processed for immune cell staining as described above. The airway localized immune cell populations were identified based on the CD45.2 antibody label. Flow cytometric analysis was performed on a Cytek Aurora and data was analyzed with FlowJo software (Tree Star). The gating strategies are shown in **Figure S3**.

### T cell depletion and IL-17 blocking experiments

For T cell depletion experiment, 8-10-week-old sex-matched mice groups received antibodies against either anti-rat isotype control or anti CD4^+^ and CD8^+^ neutralizing antibody. Mice were injected with either 500 µg anti-rat isotype antibody or a combination of 250 µg each of αCD4 and αCD8 neutralizing antibodies in PBS intraperitoneally two days (day −2) before the first HDM challenge and again on day 5 and day 12 during the ongoing HDM challenge on day 0, 7 and 14, followed by the sacrifice on day 15. Separate groups of HDM-challenged mice received 500 µg of isotype control antibodies on the same schedule to serve as controls. For IL-17 blocking, mice were injected intraperitoneally with 100µg anti-mouse anti-IL17 blocking antibody on days −1, 1, 3, 5, 6, 8, 10, 12, and 13, together with i.n. HDM challenge on day 0, day 7 and day 15, followed by sacrifice on day 15. Separate groups of HDM-challenged mice received 500 µg of isotype control antibodies on the same schedule to serve as controls. Following sacrifice, whole lung (without the post-caval lobe) is processed for flow cytometry and analysis.

### Histology experiments

Unperfused intact lung lobes were harvested from the HDM challenged mouse. Lungs were fixed in 4% paraformaldehyde for 24 hours before transferring to 1X PBS until further processing. All the lung sections were processed for paraffin embedding, microtome sectioning, and stained separately with hematoxylin and eosin (H&E) and periodic acid-schiff (PAS) stain at the Anatomy and pathology core (AMP core) at Washington University in St. Louis. Slides were imaged using the Zeiss Axioscan 7 at the Washington University Center for Cellular Imaging (WUCCI). Images were analyzed using ImageJ Fiji software as described in **Figure S4**.

### Cytokine bead array

For cytokine analysis, the post-caval lobe of the lungs was resuspended in PBS, homogenized by bead beating. Homogenates were centrifuged to isolate the cell debris-free supernatant and analyzed by BioPlex-Pro Mouse Cytokine 23-Plex Immunoassay (Bio-Rad, M60009RDPD). Assay was performed as per manufacturer’s instructions. Total protein concentration of the cell debris-free supernatant was estimated using a Bicinchoninic Acid (BCA) Assay kit (ThermoFisher, 23225). Bead array data is normalized to the total protein concentration of the individual sample and represented as picogram (pg) per milligram (mg) of total protein.

### BMDM treatment and cytokine analysis

8-12-week old sex matched mice were euthanized, followed by collection of the femur and tibia bones. Bones were flushed with complete RPMI media (Gibco, 11875135) containing 10% FBS (HiMedia, RM10432) three times to harvest the bone marrow stem cells. Red blood cells were lysed by ACK lysis buffer, followed by washing with 1X PBS to retrieve purified bone marrow cells. Bone marrow cells were differentiated in the presence of conditioned RPMI media containing the macrophage colony-stimulating factor (M-CSF), for 6 days, with additional media supplementation on day 3. After 6 days, supernatant containing the non-adherent cells was removed and adherent cells were washed with PBS three times, before collecting the differentiated macrophages for the experiment. Cultured bone marrow derived macrophages (BMDMs) were seeded in 6 well plate and treated with either mock or with 200 µg/ml HDM in RPMI media for 24 hours. Post incubation, cell supernatant was collected, centrifuged, and filtered to remove any cellular components. The cell-free supernatant was used to analyze G-CSF and KC by ELISA (R&D systems, DY414-05 for G-CSF and DY453-05 for KC) as per manufacturer’s guidelines.

### Statistical analysis for biological experiments

All the data from mouse experiments is from at least two independent experiments, unless specified otherwise. Samples represent biological (not technical) replicates of mice randomly sorted into each experimental group. No blinding was performed during animal experiments. If presented on a log scale, the data was log transformed before statistical analysis. Statistical differences were calculated using Prism 9.0 (GraphPad Software) using either unpaired Welch’s t-tests or one-way ANOVA or two-way ANOVA test with or without corrections as denoted in the figure legends. Statistically significant differences are shown for comparisons within a genotype or within treatment groups. Sample sizes were sufficient to detect differences as small as 10% using the statistical methods described. When used, center values and error bars represent means ± S.E.M. P < 0.05 was considered significant. P < 0.05 was denoted *, ** for P < 0.01, *** P < 0.001, and **** P < 0.0001. Statistical test details aad outcomes for each comparison for all graphs are available in Supplementary Dataset 1.

## AUTHOR CONTRIBUTIONS

N.D. and C.L.S. designed the project, analyzed the data, and wrote the manuscript. N.D., S.V.H., S.R.M., N.R.E., R.L.K., S.K.N., M. S., and R.W. performed the experiments. N.D., A.R., A.L.K, and C.L.S. designed the asthma protocols. A.S. assisted with the flow cytometry experiments and analysis. D.K. maintained and bred the mouse lines. A.L.K. and C.L.S. advised project design. All authors read and edited the manuscript.

## ACKNOWLEDGEMENTS

This work was supported by NIH grants R01 AI132697, U19 AI142784, Burroughs Wellcome Fund Investigators in the Pathogenesis of Infectious Disease, and the Philip and Sima Needleman Center for Autophagy Therapeutics and Research to C.L.S. N.D. is supported by Alexander and Gertrude Berg fellowship and S.K.N. is supported by a Stephen I. Morse Fellowship from the Department of Molecular Microbiology at Washington University in St. Louis School of Medicine. S.V.H. is supported by NIH grant T32 GM007067. R.L.K. is supported by a Potts Memorial Foundation postdoctoral fellowship. This material is based upon work supported by the National Science Foundation Graduate Research Fellowship Program under Grant #DGE 2139839 to N.R.E. Flow cytometry was performed at the cWIDR and Department of Molecular Microbiology Flow Cytometry Facility at Washington University School of Medicine. Histology slides were imaged at the Washington University Center for Cellular Imaging (WUCCI), which is supported by Washington University School of Medicine, the Children’s Discovery Institute of Washington University and St. Louis Children’s Hospital (CDI-CORE-2015-505 and CDI-CORE-2019-813), and the Foundation for Barnes-Jewish Hospital (3770). Histology sections were processed and stained at the Anatomy and Pathology Core (AMP core) at Washington University in St. Louis, School of Medicine. Any opinions, findings, and conclusions or recommendations expressed in this material are those of the author(s) and do not necessarily reflect the views of the National Science Foundation.

