## Supplemental figures for "Autophagy functions in lung macrophages and dendritic cells to regulate allergen-dependent inflammatory responses"

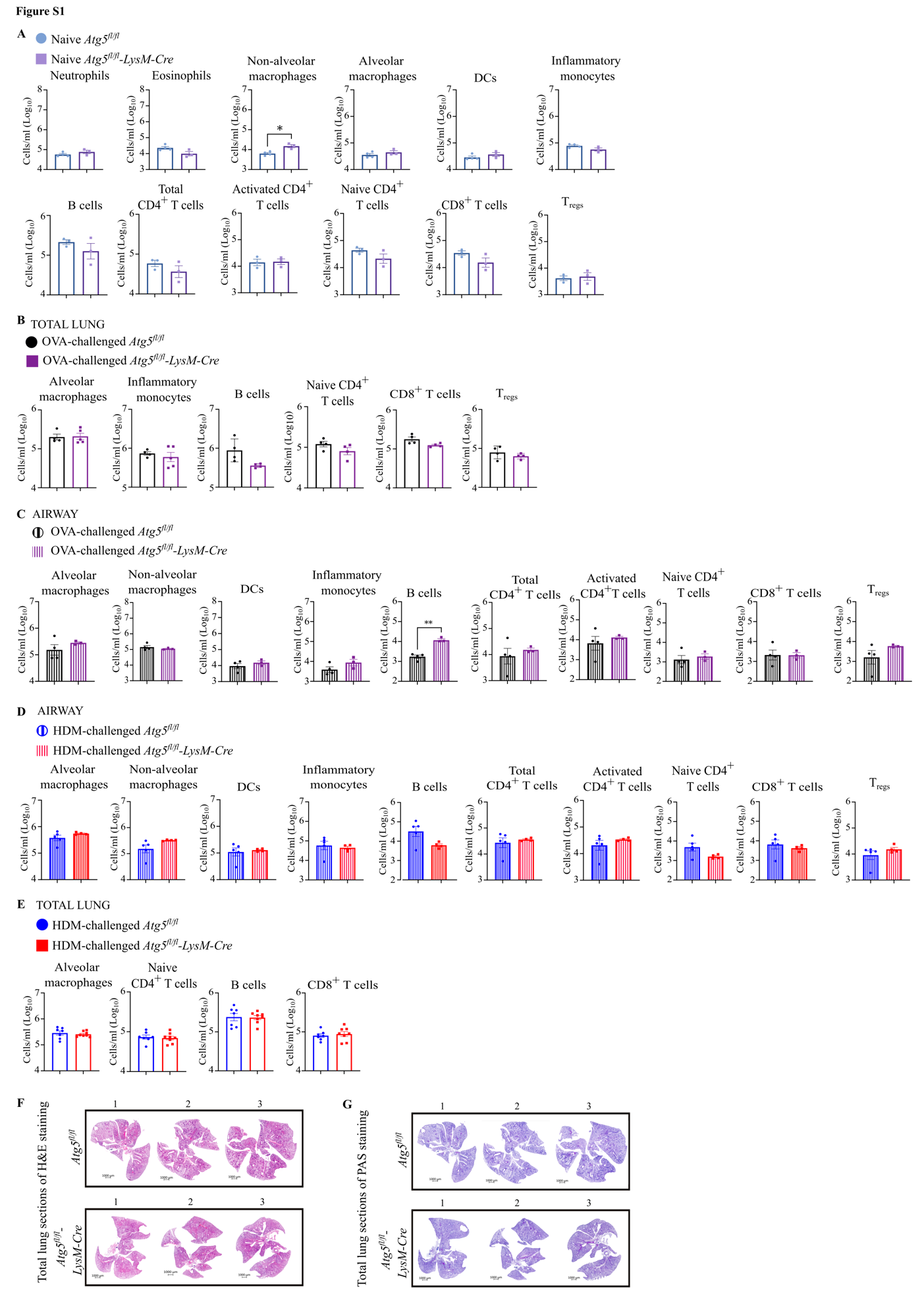

**Figure S1. Supplemental panels for Figure 1.** (**A**) Number of neutrophils, eosinophils, non-alveolar macrophages, alveolar macrophages, DCs, inflammatory monocytes, B cells, total CD4^+^ T cells, activated CD4^+^ T cells, naïve CD4^+^ T cells, CD8^+^ T cells, and T_regs_ in the lungs of naive *Atg5^fl/fl^-LysM-Cre and Atg5^fl/fl^* mice, represented as cells per ml of the lung suspension. (**B**) Number of alveolar macrophages, inflammatory monocytes, B cells, naïve CD4^+^ T cells, CD8^+^ T cells, and T_regs_ in the lungs of OVA-challenged *Atg5^fl/fl^-LysM-Cre* and *Atg5^fl/fl^* mice, represented as cells per ml of the lung suspension. (**C**) Number of alveolar macrophages, non-alveolar macrophages, DCs, inflammatory monocytes, B cells, total CD4^+^ T cells, activated CD4^+^ T cells, naïve CD4^+^ T cells, CD8^+^ T cells, and T_regs_ in the airways of OVA-challenged *Atg5^fl/fl^-LysM-Cre* and *Atg5^fl/fl^* mice, represented as cells per ml of the lung suspension. (**D**) Number of alveolar macrophages, non-alveolar macrophages, DCs, inflammatory monocytes, B cells, total CD4^+^ T cells, activated CD4^+^ T cells, naïve CD4^+^ T cells, CD8^+^ T cells, and T_regs_ in the airways of HDM-challenged *Atg5^fl/fl^-LysM-Cre and Atg5^fl/fl^* mice, represented as cells per ml of the lung suspension. (**E**) Numbers of alveolar macrophages, naïve CD4^+^ T cells, B cells, and CD8^+^ T cells in the lungs of HDM-challenged *Atg5^fl/fl^-LysM-Cre* and *Atg5^fl/fl^* mice, represented as cells per ml of the lung suspension. (**F**) H&E stained lung sections from HDM-challenge in *Atg5^fl/fl^-LysM-Cre* and *Atg5^fl/fl^* mice at 10X zoom. (**G)** PAS-stained lung sections from HDM-challenge in *Atg5^fl/fl^-LysM-Cre* and *Atg5^fl/fl^* mice. **(A-E)** Each datapoint is from an individual mouse. Data is presented on a log_10_ transformed scale. The bar graphs represent the mean +/- S.E.M. Unpaired Welch’s t-test was used to determine the statistical significance of differences, and only the statistically significant differences are shown in the graphs. * P < 0.05 and ** P < 0.01.

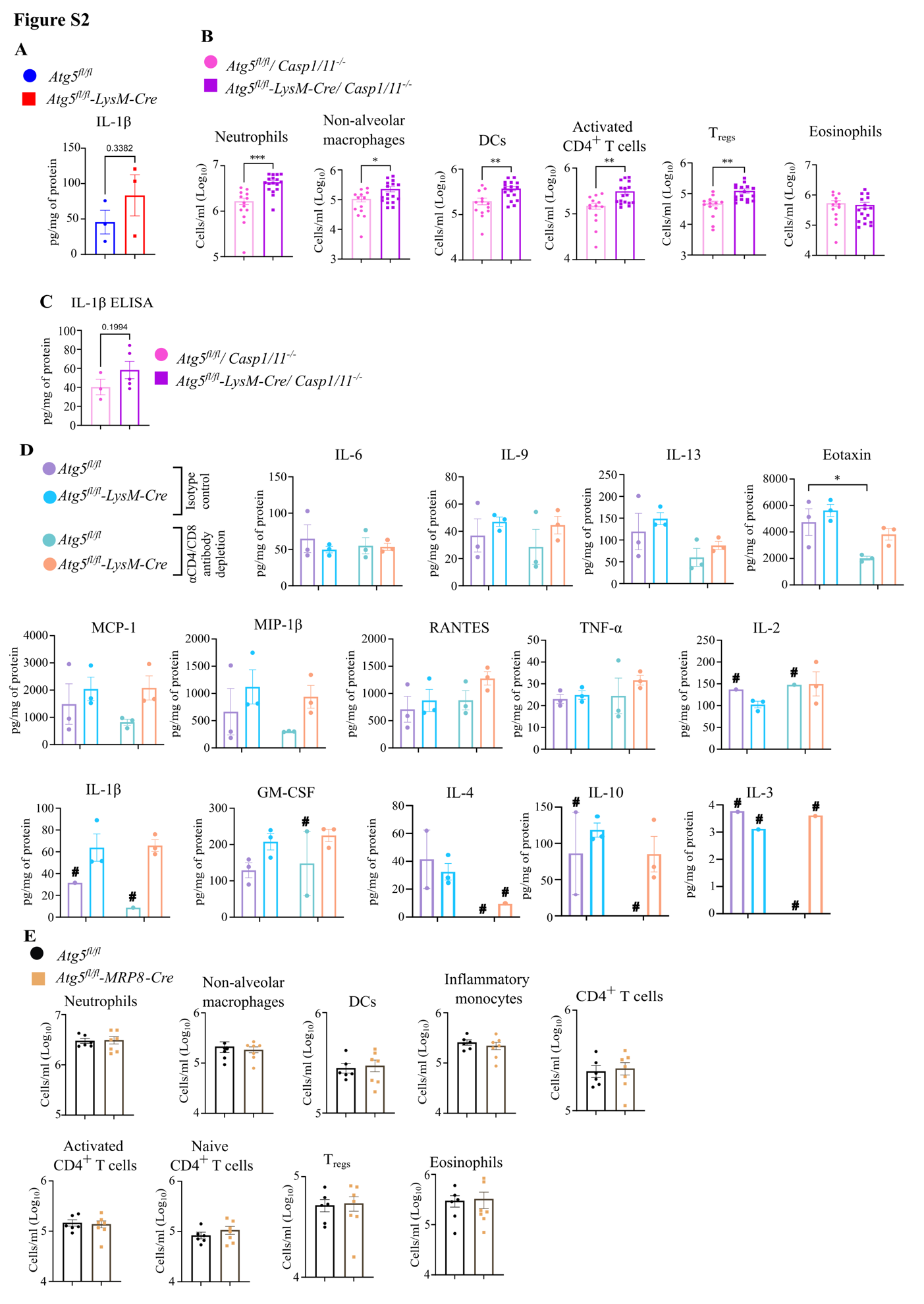

**Figure S2. Supplemental panels for Figure 4.** (**A**) Concentration of IL-1β in lung homogenate from HDM-challenged *Atg5^fl/fl^-LysM-Cre* and *Atg5^fl/^*^fl^ mice. **(B)** Number of neutrophils, non-alveolar macrophages, DCs, activated CD4^+^ T cells, T_regs_, and eosinophils in the lungs of HDM-challenged *Atg5^fl/fl^-LysM-Cre/Casp1/11^-/-^* and *Atg5^fl/fl^/Casp1/11^-/-^*mice, represented as cells per ml of the lung suspension. **(C)** Concentration of IL-1β in the lung homogenate from HDM-challenged *Atg5^fl/fl^-LysM-Cre/Casp1/11^-/-^* and *Atg5^fl/fl^/Casp1/11^-/-^*mice. **(D)** Concentration of cytokines and chemokines in total lung homogenates from HDM-challenged *Atg5^fl/fl^-LysM-Cre* and *Atg5^fl/^*^fl^ mice that received either isotype control or αCD4 and αCD8 antibodies, based on cytokine bead array. Only the data that is above the limit of detection is plotted, where # designates conditions with samples below the limit of detection. **(E)** Number of neutrophils, non-alveolar macrophages, DCs, inflammatory monocytes, total CD4^+^ T cells, activated CD4^+^ T cells, naïve CD4^+^ T cells, T_regs_, and eosinophils in the lungs of HDM-challenged *Atg5^fl/fl^-MRP8-Cre* and *Atg5^fl/fl^* mice. Each datapoint is from an individual mouse. **(B,E)** Data is presented on a log_10_ transformed scale. The bar graphs represent the mean +/- S.E.M. Unpaired Welch’s t-test was used to determine the statistical significance of differences between the groups for panels A-C, and E. Two-way Anova with Fisher’s LSD test was used to determine the statistical significance of differences between the groups in panel D. Only the statistically significant differences are shown in the graphs. * P < 0.05, ** P < 0.01, *** P < 0.001.

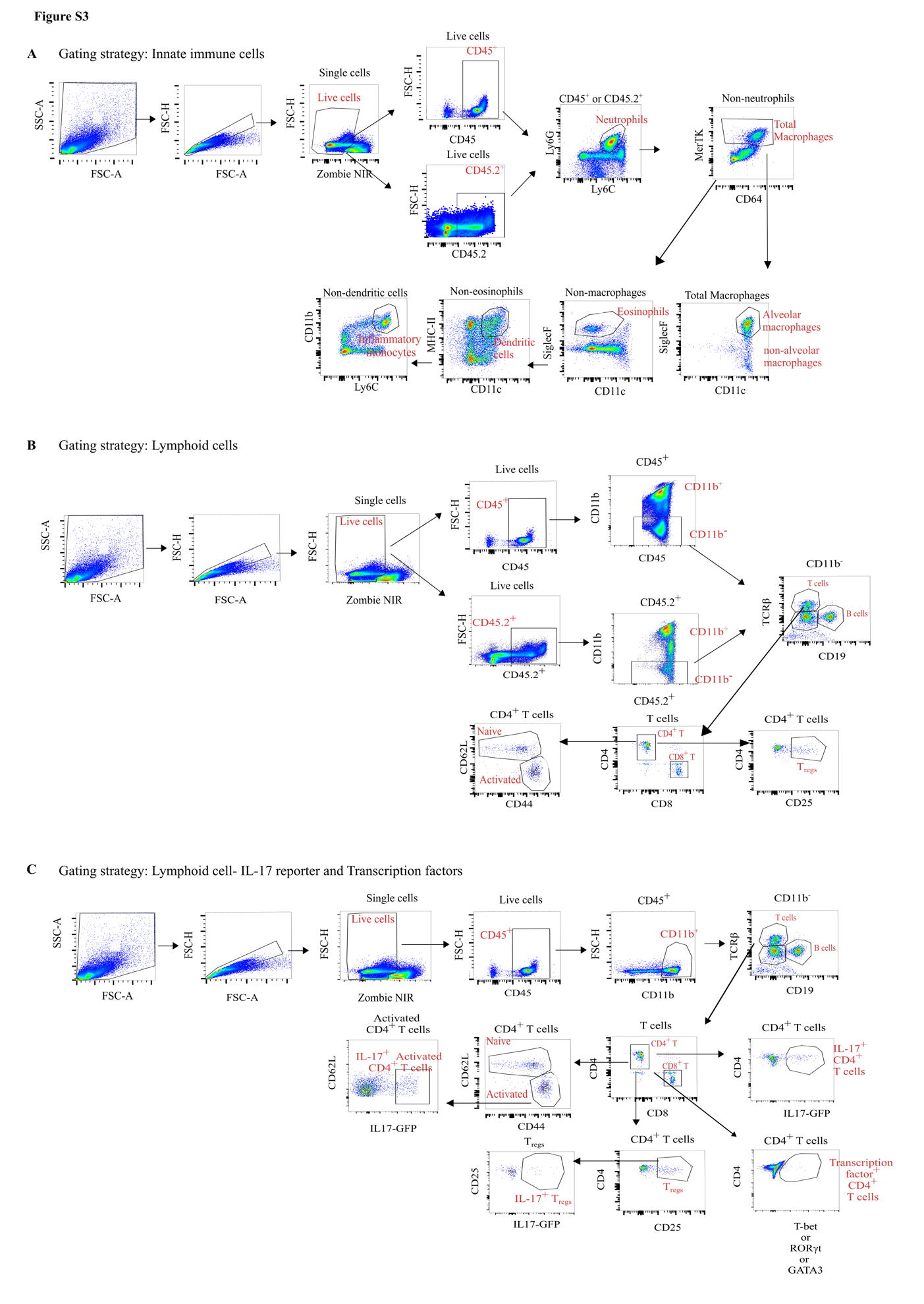

**Figure S3.** **Representative flow cytometry gating strategy.** **(A)** innate immune cells (total lung and airway specific cells) and **(B)** lymphoid cell populations (total lung and airway specific cells), and (**C**) lymphoid cell populations from total lungs that were also stained for transcription factor expression or used to detect the IL-17 GFP reporter.

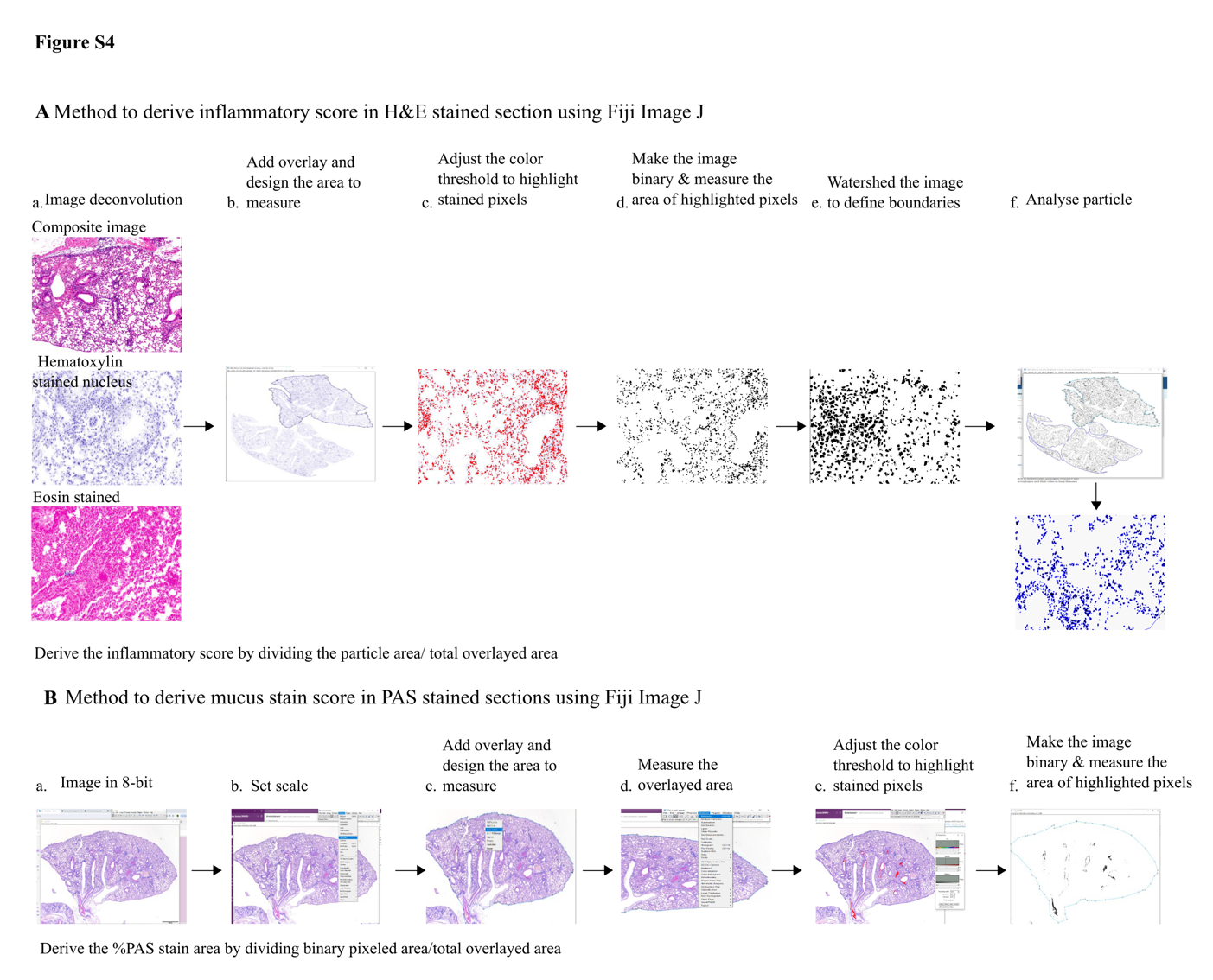

**Figure S4. Quantification methods using ImageJ. (A)** Method of quantification for the inflammatory score calculated from the H&E-stained lung sections, analyzed in Fiji ImageJ software. (**B)** Method of quantification for the mucus stain score calculated from the PAS-stained lung sections, analyzed in Fiji ImageJ software.
